# Sensory neurostimulation promotes stress resilience with frequency-specificity

**DOI:** 10.1101/2025.07.12.664223

**Authors:** Tina C. Franklin, Sara Bitarafan, Alexia T. King, Matthew Goodson, Catherine Rutledge, Tarini Gajelli, Ashley Prichard, Hector Zepeda, Nancy Kye, Steven A. Sloan, Levi Wood, Annabelle C. Singer

## Abstract

Chronic stress is a major risk factor for neuropsychiatric disorders, acting via increased neuroinflammation and disrupted synaptic plasticity. While non-invasive visual or audiovisual neurostimulation (AV flicker) at 40Hz has been shown to modulate brain immune signaling and improve cognitive performance in mouse models of Alzheimer’s disease, its effects in the context of stress remain unknown. Here we show that AV flicker protects against stress-induced behavioral, microglial, astrocytic, and synaptic changes in a sex- and frequency-specific manner. Male and female mice underwent 28 days of chronic unpredictable stress with concomitant daily AV flicker exposure at 10Hz, 20Hz, or 40Hz. Stress-induced behaviors were most effectively mitigated by 10Hz AV flicker in males and 40Hz AV flicker in females. In the medial prefrontal cortex, AV flicker normalized the balance of mature and immature dendritic spines and counteracted stress-induced molecular changes in neurons, microglia, and astrocytes, including in key neuropsychiatric risk genes. These findings show that frequency optimized AV flicker induces resilience to chronic stress.

## INTRODUCTION

Chronic or severe stress increases the risk for developing neuropsychiatric and neurodegenerative disorders by 2-fold or more^1–3^. Chronic stress causes loss of synaptic integrity, one of the most reliable predictors of neuropsychiatric symptoms and cognitive decline^4–6^. Neuroimmune signaling, microglial activation, and astrocyte reactivity are critical mechanisms of stress-induced synaptic dysfunction,^7–10^ emphasizing the need for immune-modulating interventions to counteract stress effects. We have recently shown that non-invasive flickering audiovisual (AV) neurostimulation (aka flicker) at 20Hz (within the beta frequency band) and 40Hz (within the gamma frequency band) produce distinct, frequency-specific effects on neuroimmune signaling in healthy male mice^11,12^. Furthermore, our team and others have shown that 40Hz flicker modulates microglia and astrocytes to ultimately preserve synaptic health in mouse models of neurodegeneration^13–16^. However, little is known about how flicker affects neuropsychiatric-like behavior deficits and maladaptive neuroimmune function in stressed populations. Furthermore, whether flicker can prophylactically promote resilience to stress prior to disease onset is unknown.

Chronic stress compromises synaptic integrity in the prefrontal cortex, a brain region that is highly sensitive to psychological stress, through indirect and direct mechanisms that lead to synaptic remodeling. Stress indirectly influences synaptic integrity through the activation and upregulation of proinflammatory mediators, such as pattern recognition receptors like TLR4^17^, as well as inflammatory factors like Serpina3, IL6, and CXCL2, respectively^6,17,18^. Stress also directly heightens neuronal responses that influence synaptic spines via microglia^9,19–21^ and astrocyte remodeling^8,22^. Stress alters synaptic proteins important for synaptic scaffolding and neurotransmission proteins^7,23,44^, and leads to increased glia-neuron interaction and glial synapse engulfment^6,19^. While AV flicker has been shown to affect microglia and astrocytes in healthy brains and in the context of Alzheimer’s disease pathology, it is unclear if flicker stimulation mitigates stress-induced glia changes.

Human and rodent studies show that individuals exposed to chronic or severe stress exhibit different stress susceptibility or resilience, i.e., different susceptibility to undesirable functional or physiological effects of stress. These differences in stress susceptibility are linked to the expression of specific risk and resilience genes^27,28^. Resilience can entail returning to a pre-stress state or adopting an alternative, but adaptive, beneficial response. Interventions such as exercise reduce inflammation, modify stress susceptibility, and enhance stress resilience,^29–33^ but adherence rates are low^34–36^. The potential of non-invasive brain stimulation to induce stress resilience as an alternative approach has not been widely explored. Because flicker has been shown to alter neuroimmune cells that play key roles in stress susceptibility, we wondered whether flicker at specific frequencies enhances stress resilience and promotes the expression of genes associated with stress resilience.

Given the connection between dysfunctional neuroimmune signaling, altered expression of stress-associated genes, and compromised synaptic integrity^6,28,37,38^, interventions that protect against stress-induced neuroimmune changes are of interest for at-risk populations. Because different frequencies of flicker differentially regulate neuroimmune signals and microglia^11,12^, we theorized that stimulation at an optimized frequency would protect against stress-induced neuroimmune changes and stress-induced anxiety- and depression-like behaviors. We tested the effects of 20Hz and 40Hz AV flicker because we previously found that these stimulation frequencies differentially affect neuroimmune signaling, like cytokine expression, and microglia phenotype^11,12^. We also tested the effects of 10Hz AV flicker (within the alpha band) because prior studies have shown beneficial effects of 10Hz transcranial magnetic stimulation in depression symptoms^39,40^. We used a well-established mouse model of chronic unpredictable stress (CUS)^41^ and concomitantly exposed both male and female mice to daily AV flicker intervention (10Hz, 20Hz, 40Hz, or no stimulation control) for 28 consecutive days (1hr/day). Stressed mice were subsequently screened for behavioral, cellular, and cell-specific molecular indicators of stress susceptibility or resilience following AV flicker intervention or no intervention. Specifically, we examined the effects of AV flicker on spine morphology and the transcriptomic profiles of cortical neurons, astrocytes, and microglia in stressed mice.

We were surprised to find that after 28 days of concomitant CUS and 10Hz AV flicker, males were stress resilient, behaving like unstressed control animals. Male mice exhibited protective effects of flicker in anxiety-like measures, such as the elevated plus maze test, and showed more pronounced improvements in exploratory and anxiety-like behaviors in the open field test. We found that flicker stimulation that induced stress-resilient behavior also mitigated stress-induced alterations in spine morphology, restoring proportions of mature (stubby, mushroom) to immature (long thin, filopodia) spines to levels comparable to no stress controls. Furthermore, we established that flicker stimulation that induced stress-resilient behavior counteracted stress-induced changes in gene transcription in neurons, microglia, and astrocytes. 10Hz AV flicker in male mice modulated key genes involved in stress response and regulation in neurons, astrocytes, and microglia, as indicated by differential expression of the norepinephrine receptor gene *Adra2c* in neurons, the glucocorticoid receptor regulator *Fkbp5* in astrocytes, and the glucocorticoid receptor gene *Nr3c1* in microglia. Moreover, 10Hz AV flicker mitigated stress-induced alterations in genes associated with synaptic remodeling and reactivity in neurons, astrocytes, and microglia, promoting a homeostatic state conducive to synaptic plasticity and maintenance. In females, 40Hz AV flicker was most effective at inducing behavioral resilience and coincided with synaptic and transcriptomic changes, although the effects were less pronounced. Overall, our study demonstrates that frequency-optimized flicker intervention promotes behavioral improvement, synaptic stability, and neuroimmune regulation in a cell- and sex-specific manner. These results point to a novel, non-invasive approach to prophylactically mitigate stress-induced neuropsychiatric decline.

## RESULTS

### Flicker Mitigates Stress-Induced Susceptibility to Depressive-Like and Anxiety-Like Behaviors

Chronic stress induces behavioral changes across species, including humans, non-human primates, and rodents, often presenting as depressive- and anxiety-like behaviors with individual variability. To test the effects of flicker on chronic stress, we used the well-established and validated chronic unpredictable stress (CUS) model^42^ for its ability to mimic the unpredictable and variable nature of stressors experienced in human life and to induce a range of behavioral, physiological, and neuroendocrine effects that closely resemble symptoms of human neuropsychiatric disorders such as depression. Using the CUS paradigm, we replicated stress-induced maladaptive behavioral effects in both male and female C57BL/6J mice. Mice underwent four weeks of CUS, consisting of two stressors per day (**Supplemental Figure 1**). At the end of the 4-week CUS exposure, we measured body weight change from baseline, anhedonia (via the sucrose consumption test), despair (via the forced swim test), anxiety-like behaviors (via the elevated plus maze and open field test), as well as general exploration, locomotion, and recognition memory (via the novel object recognition test) (**Figure 1A**). To capture the complexity of stress susceptibility and to quantify the overall impact of CUS, we employed an adapted version of the composite behavioral and physiological score developed by Johnson et. al (2021)^43^ (see Methods). We found a robust increase in stress-induced maladaptive physiological and behavioral changes in both sexes (**Figure 1B-C,** Males F (4, 27) = 42.54; p<0.0001; **Figure 1D-E,** Females F (4, 31) = 10.73; p<0.0001, see **Supplemental Table 11** for statistical details). CUS caused a decrease in composite stress susceptibility scores by approximately 4 standard deviations in males and 1 standard deviation in females compared to no stress controls, indicating increased stress susceptibility (**Figure 1B, Figure 1D**). The degree of these changes per behavioral assay varied by sex (**Supplemental Figure 2A-M**). To identify the primary drivers of stress susceptible compared to stress-naïve animals in both sexes, we further examined individual behavioral and physiological measures. Our results revealed that CUS induced distinct anxiety-like behaviors in both males and females. CUS exposed male mice showed significantly less center time in the open field, indicating increased anxiety-like behavior. This represented a 6.75-fold decrease on average compared to controls (**Supplemental Figure 2A**, F (4, 27) = 553.5; p<0.0001). CUS-exposed females showed increased anxiety, spending 17.4% less time on average in the elevated plus maze open arms than controls (**Supplemental Figure 2K**, F (4, 31) = 7.861; p=0.0002). These findings establish that our CUS paradigm and stress susceptible-resilience assays capture behavioral and physiological responses to chronic stress, even when those responses vary per sex.

**Figure 1:**
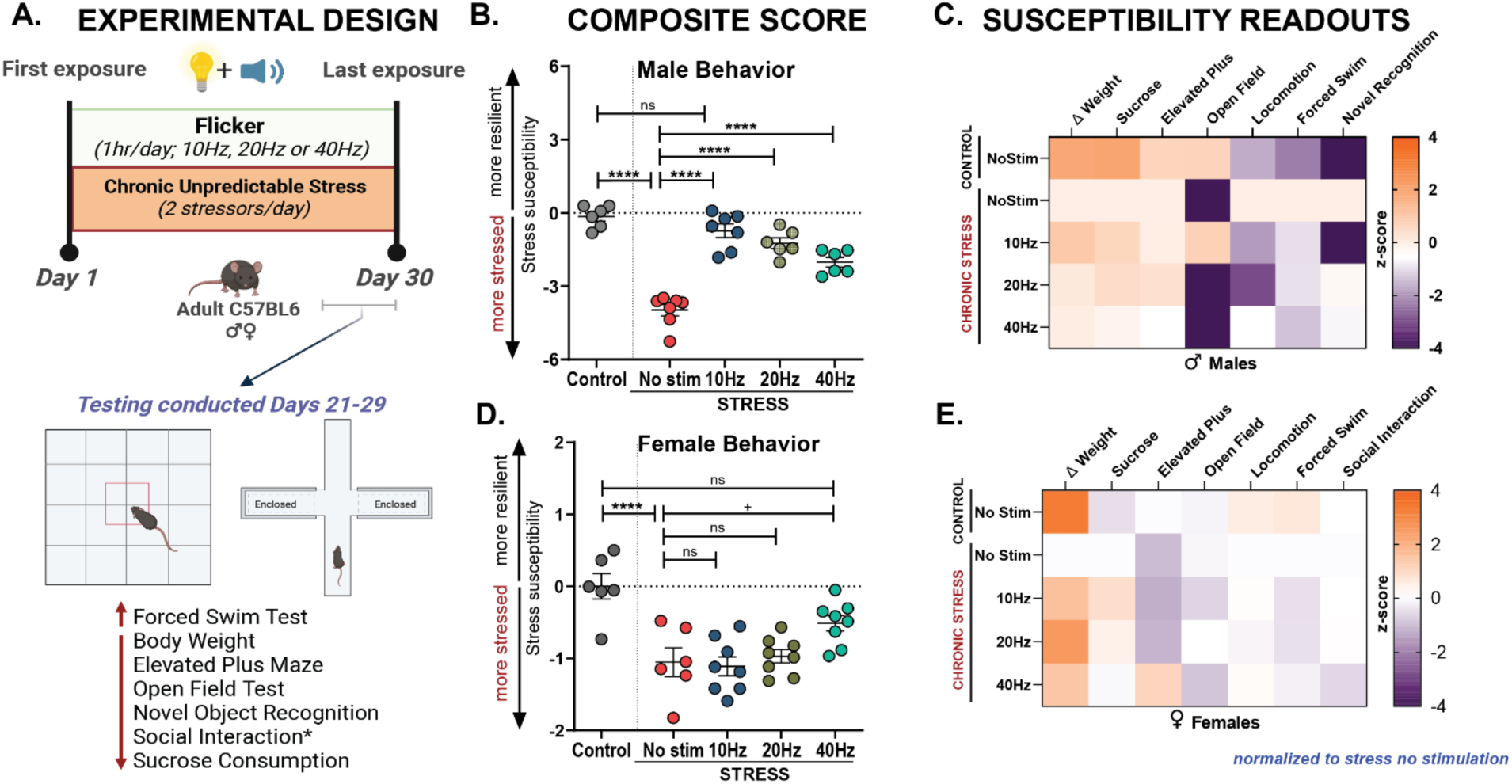
Chronic Audiovisual (AV) Flicker boosts resilience to stress in male and female mice. **(A)** Male and female mice were exposed to flicker intervention or no stimulation concomitantly with daily stress exposure. Flicker stimulation was presented at 10Hz, 20Hz, or 40Hz. Control animals (grey) underwent no stimulation and no stress exposure. Below: mice underwent a battery of behavioral assays in the last week of stress exposure. Chronic stress typically leads to decreased scores in body weight, elevated plus maze, novel object recognition social interaction and sucrose consumption and increased scores in the forced swim test. **(B)** Stress susceptibility scores generated by behavioral composite scores for male mice after stress and AV flicker intervention at 10Hz (blue), 20Hz (brown), or 40Hz (green), stress alone (red), or no stress no stimulation control (grey) conditions (F (4, 27) = 42.54; ****p<0.0001). **(C)** Matrices showing changes in body weight and behavioral performance for each assay (columns) following AV flicker intervention or stress alone normalized to stress no stimulation. Color indicates increased (orange) or decreased (purple) activity of specific behaviors (z-scored). For body weight, sucrose consumption and elevated plus maze (first four columns), stress led to lower z-scores as expected, while flicker increased these scores. For locomotion, forced swim, and novel recognition tests (last 3 columns), stress results in higher z-scores as expected while flicker counteracted these effects. **(D)** As in B for females (F (4, 31) = 10.73, ****p<0.0001). **(E)** As in C for females. B and D show mean ± SEM; n = 6-8 animals per group. +p=0.0525, ****p<0.0001, ns, not significant. One-way analysis of variance performed for compiled scores followed by Bonferroni correction for multiple comparisons. Experimental paradigm graphics created in BioRender. Franklin, T. (2025) https://BioRender.com/h87x345. See **Supplemental Table 11** for statistical details.

To investigate whether AV flicker mitigates stress-induced physiological and behavioral dysregulation, we exposed a subset of CUS-exposed mice to daily AV flicker at 10Hz, 20Hz, or 40Hz frequencies on the same day but at a different time than CUS exposure (**Figure 1A, Supplemental Figure 1**). We found that that AV flicker significantly alleviated stress-induced behavioral deficits in a frequency-dependent manner. Specifically, AV flicker mitigated depressive-like and anxiety-like behaviors in both sexes, with the most pronounced effects resulting from 10Hz AV flicker exposure in males and 40Hz AV flicker exposure in females. In males, 10Hz AV flicker improved stress susceptibility scores, marked by an improvement of approximately 3 standard deviations (**Figure 1B,** Males F (4, 27) = 42.54; p<0.0001) while in females, 40Hz AV flicker resulted in approximately 1 standard deviation improvement (**Figure 1D,** Females F (4, 31) = 10.73; p=0.0001) relative to unstimulated stress groups. These findings show that AV flicker promotes resilience against stress-related physiological and behavioral vulnerabilities.

Given that we observed that chronic stress effects were most pronounced in male mice and flicker produced the strongest behavioral response in males, we focused on the effects of flicker stimulation in stressed male subjects, while still including the effects in females. To understand the cellular and molecular changes related to flicker-induced resilience, we further assessed the optimal frequency of flicker in each sex that most effectively induced resilience, namely 10Hz in males and 40Hz in females.

### Flicker Mitigates Stress-Induced Synaptic Remodeling

Because chronic stress is known to induce neuropsychiatric decline through its disruptive effects on synaptic health and plasticity, we next asked how CUS and AV flicker, separately and together, affected neuronal spines in the medial prefrontal cortex (mPFC). mPFC undergoes substantial neuroplastic changes under chronic stress, including dendritic retraction, spine loss and alterations in spine morphology, collectively referred to as spine dynamics^44–47^. We hypothesized that AV flicker frequencies that mitigated stress-induced-behaviors would also prevent synaptic loss and maladaptive spine dynamics. To assess the effects of AV flicker on synaptic integrity, we used Thy1-EGFP reporter mice, which express enhanced green fluorescent protein (EGFP) in cortical neurons to allow for visualization of dendritic spines (**Figure 2A-B**). Mice were subjected to CUS for 3 weeks, followed by daily flicker exposure (1 hour/day) at either 10Hz or 40Hz, frequencies shown to be most effective at mitigating stress-induced behavioral changes in males or females, respectively (**Figure 2A-B, Supplemental Figure 2A-B**).

**Figure 2:**
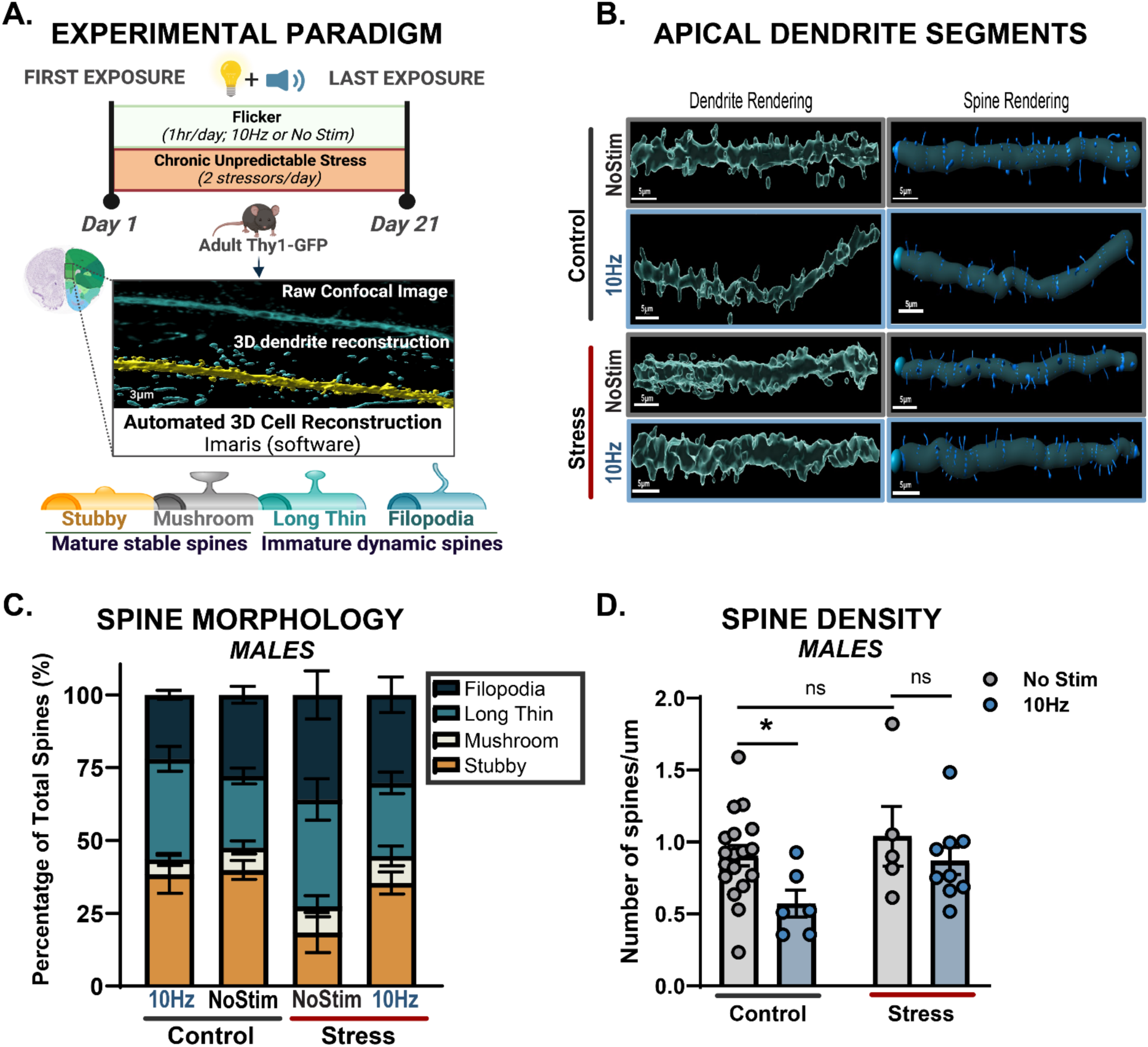
Chronic AV flicker maintains ratio of dynamic and stable cortical dendritic spines of stressed mice. **(A)** Top: Experimental paradigm. Bottom: Super-resolution confocal images and 3D morphology reconstruction of apical dendritic spines in Imaris imaging software software identified mature stable spines including stubby (orange) and mushroom (grey), and immature dynamic spines including long thin (light turquoise) and filopodia (dark turquoise). **(B)** Representative images of targeted apical dendrite segments from layer II/III of medial prefrontal cortex showing dendrite (left) and spine (right) rendering. (**C**) Spine morphology of each type in male mice following either no flicker (“NoStim”) or 10Hz AV flicker (“10Hz”) in no stressed (“Control”) and CUS (“Stress”) conditions (F (9, 136) = 2.498, p=0.0112). 10Hz AV flicker normalizes immature to mature spine ratio in stressed males exposed compared to stress no stimulation group (F (1, 34) = 4.250; p=0.0470), driven by the maintenance of stubby spines (F (1, 34) = 5.872; p= 0.0209). **(D)** Spine density in male mice following either no flicker (control no stim (grey bar over grey line) and stress no stim (grey bar over red line) or AV flicker (control 10Hz (blue bar over grey line) or stress 10Hz (blue bar over red line)) in no stress and stress conditions (F (1, 34) = 4.888, p=0.0339). *p<0.05. Error bars show mean ± SEM; n = 5-18 dendrite segments from 4-7 animals per group. Each dot is a dendrite segment. Morphological analyses were conducted at the group level, with data aggregated across dendrite segments rather than averaged per subject. Two-way analysis of variance performed for compiled scores followed by Bonferroni correction for multiple comparisons in spine morphology. Two-way analysis of variance performed for compiled scores followed by uncorrected Fisher’s LSD for multiple comparisons in spine density dataset. See **Supplemental Table 11** for statistical details.

We first established that CUS affected spine types, decreasing mature and increasing immature spines. After CUS exposure, we quantified apical dendritic spine density and morphology in layer II/III of the mPFC, layers particularly vulnerable to stress-induced synaptic remodeling^44,48^. Stress exposure significantly altered the ratio of dendritic spine types in males and females, assessed separately. In males, CUS lead to a 42% reduction in mature spine types (mushroom and stubby spines) and a 38% increase in immature spines (long thin and filopodia) on average compared to no stress controls (**Figure 2C**, F (9, 136) = 2.498; p=0.0112). This effect was primarily driven by a significant reduction in the density of stubby spines (F (1, 34) = 5.872; p=0.0209) in the unstimulated stressed males. Females exhibited a different response (**Supplementary Fig 3A-C,** F (9, 428) = 4.814; p<0.0001), primarily driven by changes in density of long thin spines (F (1, 107) = 13.88; p=0.0003). These changes are indicative of disrupted synaptic stability and plasticity in the stressed groups consistent with previous reports of stress-induced changes in spine dynamics, in studies of chronic or repeated stress models^24–26^.

We found that AV flicker intervention restored typical proportions of mature and immature dendritic spine types in stressed animals (**Figure 2C,** F (9, 136) = 2.498; p=0.0112, **Supplemental Figure 3C** F (9, 428) = 4.814; p<0.0001). Notably, 10Hz AV flicker intervention in stressed males increased the number of mature (stubby and mushroom) spines by 63% on average compared to stressed groups (**Figure 2C,** F (1, 34) = 4.250; p=0.0470) while 40Hz flicker-exposed stressed females showed normalization of immature spines primarily through a 37% reduction on average in long thin spines (**Supplemental Figure 3C,** F (1, 107) = 5.010; p=0.0273). These results show that AV flicker mitigated stress-induced changes in the ratio of mature and immature spine types in stressed mice in a frequency- and sex-specific manner, suggesting that AV flicker promotes synaptic resilience by counteracting the synaptic deficits induced by chronic stress.

We found that AV flicker alone was sufficient to promote increases in spine density in no stress control males (**Figure 2D,** Males F (1, 34) = 4.888; p=0.0339) and no stress control females (**Supplemental Figure 3D,** Females F (1,107) = 3.785; p=0.0543) compared to no stress unstimulated groups. Surprisingly, the 3D dendritic reconstructions revealed no significant changes in spine density along apical dendrite segments in stress unstimulated groups compared to no stress unstimulated controls (**Figure 2D,** uncorrected Fisher’s post hoc males control no stim vs. stress no stim p=0.3947, **Supplemental Figure 3D,** uncorrected Fisher’s post hoc females control no stim vs. stress no stim p=0.4896).

### Chronic Stress Induces Cell-Type Specific Transcriptional Profile Changes in the Frontal Cortex

Because synaptic spines are collectively regulated by neurons, astrocytes, and microglia, we next examined transcriptional profiles of each of these cell types in response to CUS and different frequencies of flicker. We first established that microglia are required for behavioral and physiological responses to chronic stress (**Supplemental Figure 4A**). Previous studies have shown that modulation or depletion of microglia prevents stress-induced anxiety-like behaviors, suggesting that microglia responses to stress plays a critical role in stress susceptibility^49–54^. Interestingly, PLX3397-treated male mice showed attenuation in stress-induced responses including body weight (F (2, 19) = 11.17; p=0.0006), forced swim test (F (2, 19) = 5.230; p=0.0155), and elevated plus maze performance (F (2, 21) = 1.769; p=0.1951), indicating that microglia play a major role in mediating these stress effects (**Supplemental Figure 4B-E**). However, stress-induced changes in open field test center entries (F (2, 19) = 3.557; p=0.0487) and sucrose consumption (F (2, 21) = 9.867; p=0.0010) were maintained or exaggerated in PLX3397-treated males. These results indicated that microglia play a critical role in mediating the effects of stress on some, but not all, anxiety-like and depressive-like behaviors. Therefore, we hypothesized that in addition to microglia, neurons and astrocytes are also affected by stress because they also contribute to synaptic remodeling^8,22^. To investigate this, we assessed how stress impacts transcriptional profiles in microglia, astrocytes, and neurons. Because we wanted sufficient sensitivity to detect lowly abundant immune genes, we performed bulk RNA sequencing on purified cell populations isolated via immunopanning^55–57^. Specifically, we enriched and analyzed Thy1+ frontal cortical neuron (which is comprised of both excitatory and inhibitory neurons), CD45+ microglia, and HepaCam+ astrocyte samples isolated from adult C57BL/6J mice subjected to concomitant CUS and AV flicker stimulation at frequencies of 10Hz, 40Hz, or no stimulation (**Figure 3A**; see Methods). As described above, we focused on the frequencies of stimulation shown to best induce stress resilience in males and females.

**Figure 3:**
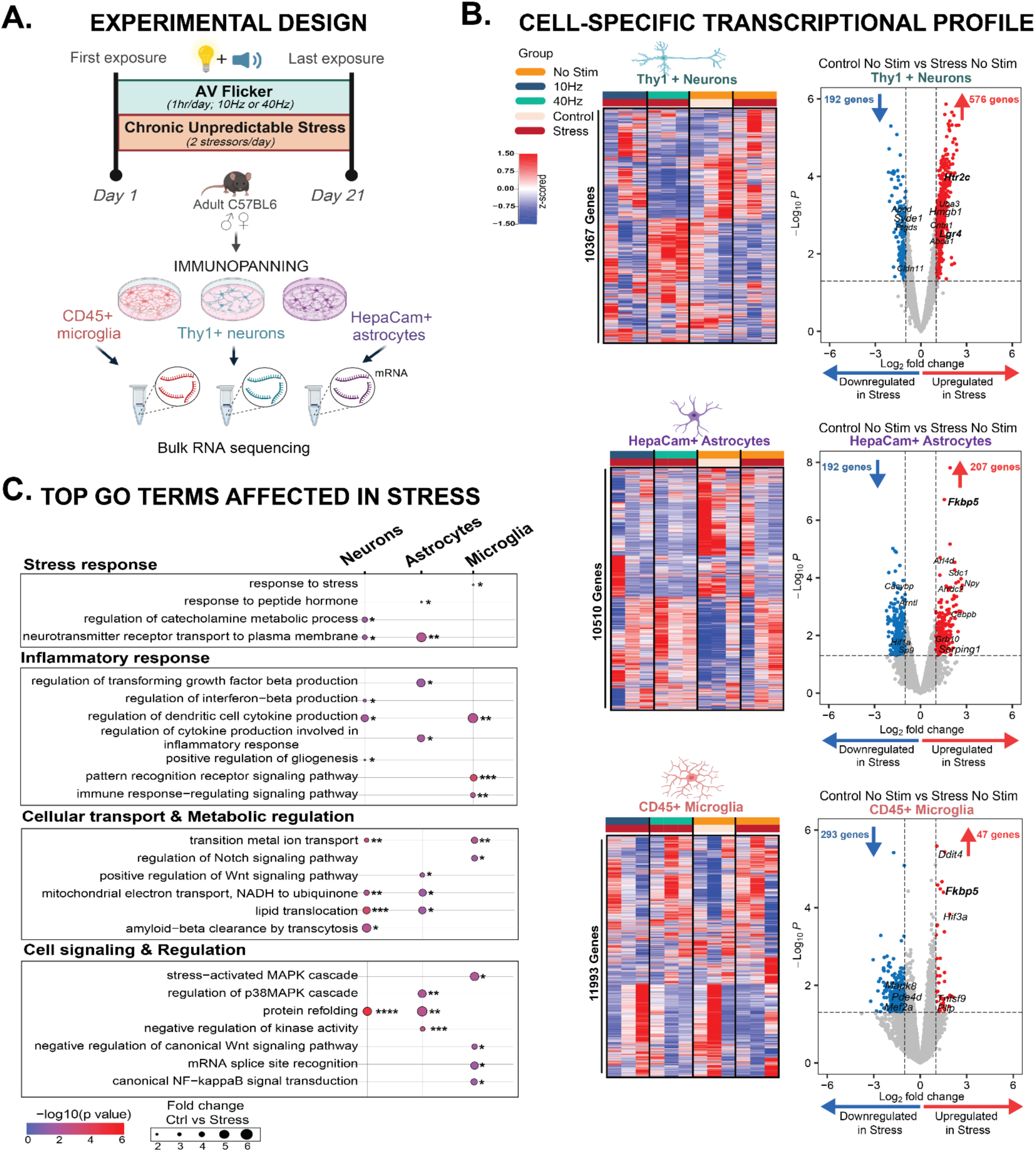
Chronic stress robustly shifts neuronal, astrocyte, and microglia transcriptome profile in males. **(A)** Experimental paradigm: Thy1+ neurons, HepaCam+ astrocytes, and CD45+ microglia were isolated by immunopanning within 1 hour after the final stressor exposure on the last day of the paradigm. Genes involved in the regulation of stress signaling are highlighted in bold. **(B)** Heatmaps of gene expression of neurons (top), astrocytes (middle), and microglia (bottom) in control no stim (orange/light pink) or following stress alone (dark red/orange), stress combined with AV flicker at 10Hz (dark red/blue), or 40Hz (dark red/green). Colors above the heatmap indicate stress (dark red) or no stress (pink) and no stimulation (orange), 10Hz (blue), or 40Hz (green) flicker exposure. In the heatmap, red shows increased expression and blue shows decreased expression compared to no stress no stimulation controls (threshold set. log2fc>1 and p<0.05). **(C)** Primary pathways affected by stress alone in neurons, astrocytes, and microglia. N = 3 pooled samples from 6 animals per group. Transcriptional profile at gene expression level unadjusted p<0.05, Wald test. GO terms pathway analysis *p < 0.05, **p<0.01, ***p<0.001, and ****p<0.0001. **A** Created in BioRender. Franklin, T. (2025) https://BioRender.com/r17l695. See **Supplemental Table 11** for statistical details.

We identified significant differential expressed genes (DEGs) across neurons, microglia, and astrocytes, revealing key insights into how stress modulates cell-type-specific signaling in the frontal cortex (which encompasses the medial prefrontal cortex) of male and female mice (**Figure 3B**, **Supplemental Figure 5B, Supplemental Figure 6, Supplemental Tables 1-3)**. DEGs were defined to be genes with fold change greater than 2 or less than -2 and unadjusted p<0.05 (DESeq2, see Methods). To gain insight into the functional implications of these DEGs, we performed an over-representative gene ontology (GO) analysis (unadjusted p<0.05, Fisher’s exact test; GO terms were unadjusted for comparing stress and no stress groups to be more inclusive, see Methods). Our analysis uncovered four major classes of GO terms affected by chronic stress in males: stress response, inflammatory response, cellular transport and metabolic regulation, and cell signaling and regulatory cascades (**Figure 3C, Supplemental Table 4**, unadjusted p<0.05; see Methods). Notably, microglia, astrocytes, and neurons all displayed significant upregulation of stress-related genes, with DEGs known to be involved in neuropsychiatric diseases^27,28,58^. In neurons, genes such as *Htr2c, Gpr88, Lgr4* and *Ptgds* were differentially expressed after stress (**Figure 3B, Supplemental Table 1**). In astrocytes, *Ier2, Nfkbia, Cdkn1a,* and *Apod* were upregulated by stress (**Figure 3B, Supplemental Table 2**). Similarly, in microglia, *Ddit4, Sgk3, Tlr4,* and *P2ry12* were differentially expressed between stress and no stress groups (**Figure 3B, Supplemental Table 3**). Importantly, the key stress regulation gene *Fkbp5* was upregulated in both astrocytes and microglia, suggesting shared stress signaling effects across both cell types (**Figure 3B, Supplemental Table 2, Supplemental Table 3**).

Of the DEGs in Thy1+ neurons, 75% were up-regulated in males and 54% were up-regulated in females (**Figure 3B, Supplemental Figure 5B**, **Supplemental Table 1**). Furthermore, neurons exhibited notable alterations in cellular transport and metabolic pathways (i.e., *Htr2c*, *Adra2c*, *Slc6a4* and *Syde1*), which are critical for synaptic function and neuronal communication. Interestingly, immunomodulatory genes (i.e., *Lgr4, Aldh1a1* and *Abca1*) were also significantly altered in neurons (**Figure 3B**, **Supplemental Figure 5B, Supplemental Table 1**), which may be consistent with glial-mediated repair in response to stress-induced damage. Thus, these genes highlight how CUS affect pathways involved in neuronal-glial communication, which may play a pivotal role in stress-induced synaptic remodeling.

Microglia showed more down-regulated DEGs following CUS (**Figure 3B**, **Supplemental Figure 5B**, **Supplemental Table 3**). Only 14% of DEGs in male microglia and a mere 6% in female microglia are upregulated. Despite the overall trend of downregulation in glial cells, the genes that were upregulated play crucial roles in inflammatory and stress responses. In microglia, stress-induced upregulation of inflammation-related genes such as *Ddit4, Fkbp5,* and *Tnfsf9* suggests increased reactivity.

Astrocytes also showed a bias towards downregulation in response to CUS, with 65% of DEG down regulated in males and 50% downregulated in females (**Figure 3B, Supplemental Figure 5B**, **Supplemental Table 2**). In astrocytes, the upregulation of genes like *Serpina3n, Cebpb, Cebpd and Fkbp5* in response to CUS indicates an enhanced reactivity state. This selective reactivity of cortical microglia and astrocytes occurs against a backdrop of overall gene downregulation, possibly indicating a complex regulatory mechanism where most cellular functions are suppressed while stress-response and inflammatory pathways are enhanced, suggesting that these cells are primed to modulate neurotoxicity, synaptic remodeling and pruning observed in the stressed mPFC. These changes show that CUS induces molecular alterations in neuron-glia interaction signaling in our subjects.

### Flicker Mitigates Stress-Induced Transcriptional Changes in Neurons

Because AV flicker mitigated stress-induced behavioral changes, we next determined if exposure to specific flicker frequencies mitigated chronic stress-induced transcriptional changes. We were especially interested in the different transcriptional effects of 10Hz and 40Hz AV flicker because, while both are flickering stimuli, 10Hz AV flicker in males resulted in the most robust behavioral stress resilience compared to 40Hz. Thus, specific transcriptional effects of 10Hz AV flicker reflect changes related to stress resilience, rather than flicker exposure broadly. To assess how frequency-specific flicker modulation affects stress-induce gene regulation, we identified DEGs that were modified by chronic stress and showed changes in the opposite direction with flicker intervention. These *’modifiable markers’* were upregulated by stress and downregulated by AV flicker, or vice versa.

In isolated Thy1+ neurons, we found that the overall number of DEGs regulated by 40Hz AV flicker was 12-fold larger than the number DEGs regulated by 10Hz in males, 2987 DEGs versus 240 DEGs respectively (**Figure 4, Supplemental Table 1**). However, 10Hz AV flicker resulted in a greater percentage of modifiable DEGs (**Supplemental Figure 7A, Supplemental Table 5**; see Methods) and mitigated key stress-induced gene expression changes (**Figure 4B, 4C**). Specifically, 23% of genes differentially regulated by 10Hz AV flicker were modifiable DEGs while only 12% of genes regulated by 40Hz AV flicker were modifiable markers (**Supplemental Figure 7A**). To assess overlapping effects of these different flicker frequencies, we identified 37 DEGs modulated by both 10Hz and 40Hz AV flicker interventions (**Figure 4C, Supplemental Figure 7C**). GO analysis of these overlapping DEGs revealed their primary involvement in transcription, mitochondria function, and response to stimulus or stress (**Figure 4C, Supplemental Figure 7D, Supplemental Table 6,** unadjusted p<0.05; see Methods). Together, these results show that while 40Hz AV flicker intervention had broader transcriptional effects in neurons, 10Hz AV flicker more specifically counteracted the transcriptional effects of stress.

**Figure 4:**
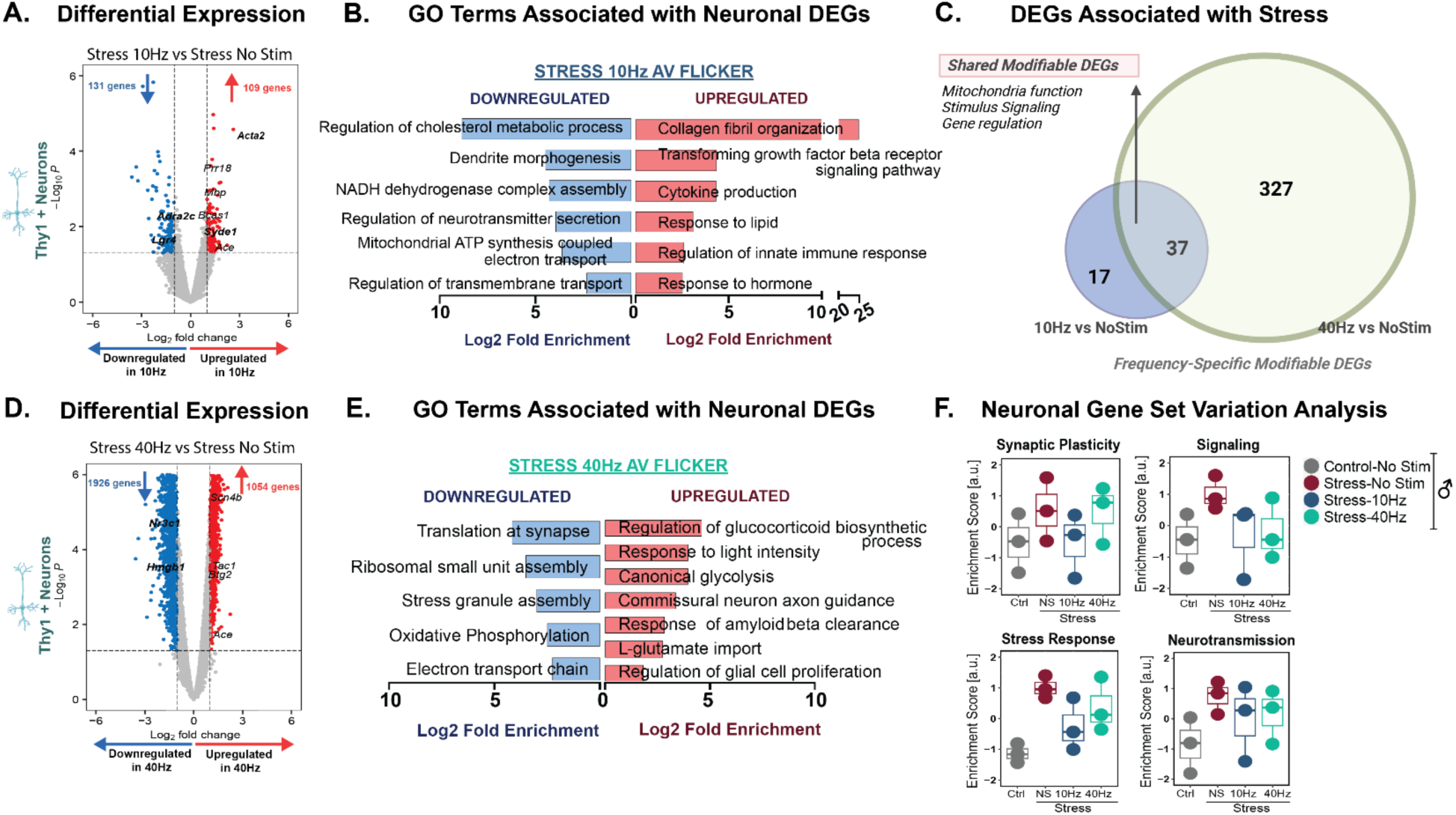
10Hz AV flicker mitigates stress-induced transcriptional changes in neurons. **(A)** Volcano plot showing differential gene expression of cortical neurons in stressed males following 10Hz AV flicker compared to stress no stimulation (threshold set. log2fc>1 and p<0.05). Genes involved in the regulation of stress signaling are highlighted in bold. **(B)** Gene ontology (GO) enrichment analysis for genes downregulated (blue) and upregulated (red) after 10Hz AV flicker exposure in stressed males compared to no stimulation stressed males. **(C)** Venn diagram demonstrating unique and shared (pink) modifiable enriched genes associated with stress following 10Hz (blue) AV flicker compared to 40Hz (green) AV flicker. **(D)** and **(E)** as in A and B for 40Hz AV flicker exposure in stressed males compared to no stimulation stressed males. **(F)** Enrichment scores for specific biological processes (i.e. stress response and synaptic signaling) in stressed male mice under different flicker exposure conditions. Box plots show the distribution of scores across samples (median ± Interquartile range; n = 3 pooled samples from 6 animals per group). Transcriptional profile at gene expression level without FDR adjustment p<0.05, Wald test. GO terms pathway analysis FDR-adjusted, p < 0.05. See **Supplemental Table 11** for statistical details.

10Hz AV flicker resulted in robust downregulation of pathways related to metabolic and synaptic function and upregulated adaptive extracellular matrix remodeling and immune response pathways, processes known to play a role in synaptic stabilization (**Figure 4B, Supplemental Table 7,** FDR-adjusted p<0.05; see Methods). While both 10Hz and 40Hz AV flicker mitigated stress induced molecular changes in males and females (**Figure 4**, **Supplemental Figure 6, Supplemental Figure 8, Supplemental Table 4**), 40Hz AV flicker induced a more moderate response than 10Hz AV flicker. While 40Hz AV flicker also downregulated key metabolic pathways such as oxidative phosphorylation and electron transport chain, the magnitude of change was lower compared to 10Hz AV flicker (**Figure 4B, 4E, Supplemental Table 7,** FDR-adjusted p<0.05; see Methods). 40Hz AV flicker-upregulated pathways included glucocorticoid regulation, canonical glycolysis, and response to light, alluding to the enhanced metabolic and neuroendocrine regulation under 40Hz AV flicker. Importantly, 40Hz AV flicker also restored pathways annotated for commissural neuron connectivity, pointing to a role for flicker in synaptic repair and plasticity. Custom gene set variance analysis (GSVA) revealed a trend of difference between 10Hz and no stimulation stress group with 10Hz more similar to no stress controls; (**Figure 4F**, not significant; see Methods)^59^. Together these results show that 10Hz AV flicker counteracts the transcriptional effects of stress with key effects on genes involved in synaptic function, metabolism, and immune response pathways. These transcriptional changes in response to 10Hz AV flicker coincided with the most improved behavioral outcomes and normalized synaptic spines in response to flicker stimulation. Thus, our transcriptomic analysis reveals how 10Hz AV flicker specifically counteracts the transcriptomic effects of stress in neurons, in comparison to 40Hz AV flicker, which has broader but less stress-specific effects. These findings highlight the distinct mechanisms by which different frequencies of AV flicker modulate stress responses at the gene expression level.

### 10Hz AV flicker Reversed the Effects of Stress on Clinically Relevant *Fkbp5*

Because astrocytes play crucial roles in synaptic remodeling and stress-induced synaptic dysfunction, we hypothesized that flicker modulates stress-induced changes in astrocyte transcription. Therefore, we next examined the transcriptomic effects of AV flicker on frontal cortical astrocytes in the context of chronic stress. We analyzed isolated HepaCam+ astrocytes and observed 47 DEGs in response to 10Hz AV flicker in stressed males (**Figure 5A, Supplemental Table 2**), while 40Hz AV flicker stimulation resulted in 404 DEGs (**Figure 5B, Supplemental Table 2**). As in our neuronal analysis, we identified modifiable DEGs that were altered by chronic stress and showed reversed expression patterns with flicker **(Supplemental Figure 7, Supplemental Table 4**). Interestingly, while more genes were differentially expressed after 40Hz AV flicker, 10Hz AV flicker was more effective at counteracting stress-induced transcriptional alterations in male mice. Specifically, 9% of genes differentially regulated by 10Hz AV flicker were modifiable markers while only 1% of genes differentially regulated by 40Hz AV flicker were modifiable markers (**Supplemental Figure 7A**) with no overlap between modifiable genes affected by 10Hz and 40Hz AV flicker (**Figure 5C**). Importantly, analysis of clinically relevant genes revealed that 10Hz AV flicker specifically modulated key neuropsychiatric risk genes altered by chronic stress, most notably reversing stress-induced upregulation of *Fkbp5* (**Figure 5A**), a well-established biomarker of stress susceptible and psychiatric disorders^60–62^.

**Figure 5:**
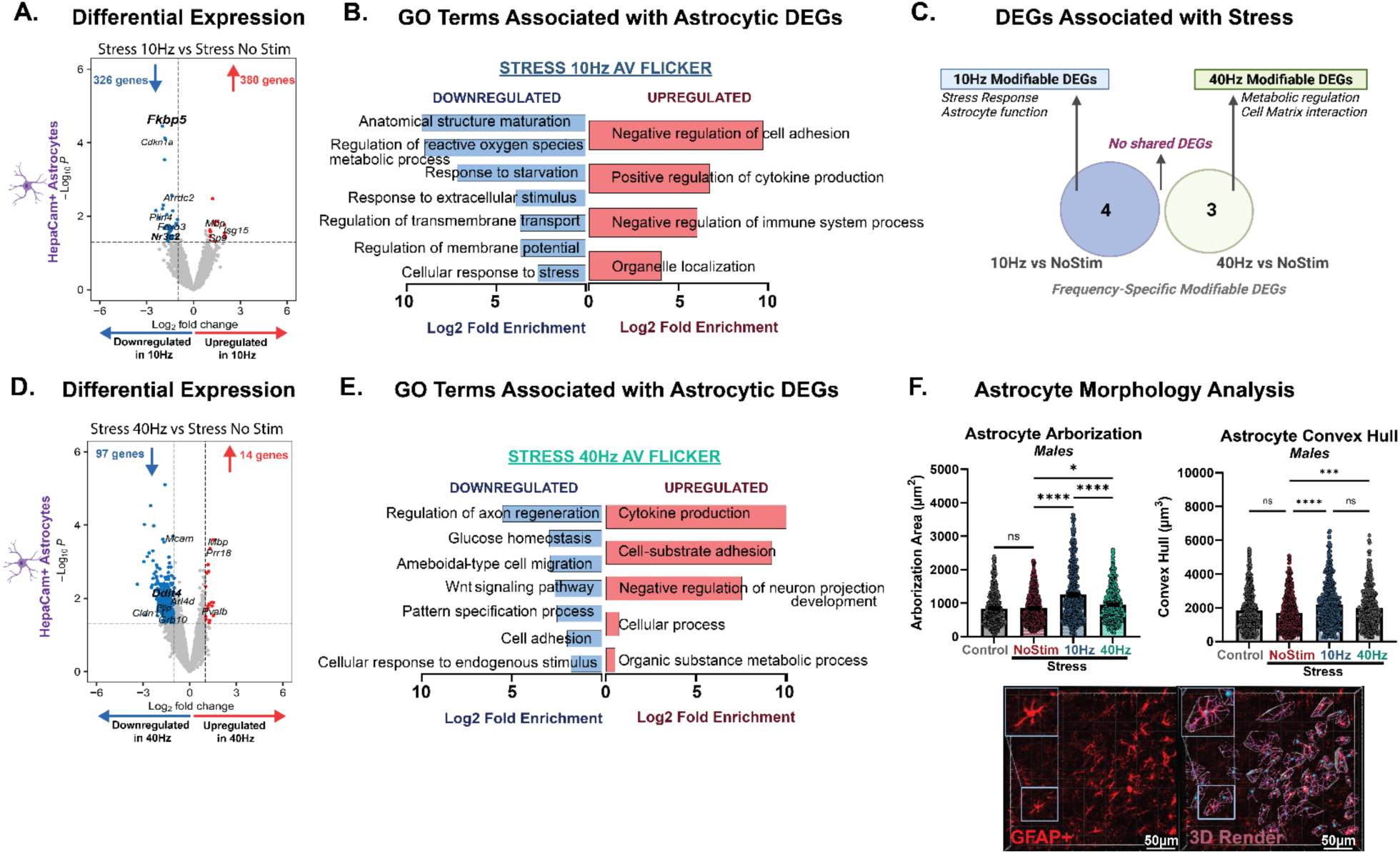
10Hz AV flicker reverses the effects of stress on clinically relevant *Fkbp5* and mediated stress-induced transcriptional changes in frontal cortical astrocytes. **(A)** Volcano plot showing differential gene expression of astrocytes in stressed males following 10Hz AV flicker compared to stress no stimulation (threshold set. log2fc>1 and p<0.05). Genes involved in the regulation of stress signaling are highlighted in bold. (**B)** Gene ontology (GO) enrichment analysis for genes downregulated (blue) and upregulated (red) after 10Hz AV flicker exposure in stressed males compared to no stimulation stressed males. **(C)** Venn diagram demonstrating unique modifiable enriched genes associated with stress following 10Hz (blue) AV flicker compared to 40Hz (green) AV flicker. **(D)** and **(E)** as in A and B for 40Hz flicker exposure in stressed males compared to no stimulation stressed males. **(F)** Cortical astrocyte branching (arborization, left) and convex hull volume (right) in stressed male mice under AV flicker intervention at 10Hz (blue), 40Hz (green), stress alone (red), or no stress no stimulation control (grey) conditions (arborization (H (3) = 84.70; p<0.0001); convex hull (H (3) = 43.38; p<0.0001)). Below: Example 20x confocal image of GFAP+ astrocytes (left) and reconstruction of morphology (right), scale bar 50µm. Error bars show mean ± SEM. Each dot is a cell. N = 3 pooled samples from 6 animals per group for transcriptome analysis and n = 439-593 cells from 5 animals per group for morphology analysis. Morphological analyses were conducted at the group level, with data aggregated across cells within a group. Transcriptional profile at gene expression level without FDR adjustment p<0.05, Wald test. GO terms pathway analysis FDR-adjusted, p<0.05. Kruskal-Wallis test performed for astrocyte morphology analysis followed by Dunn’s post hoc test. *p<0.05; ***p<0.001; ****p<0.0001. See **Supplemental Table 11** for statistical details.

Despite having fewer DEGs, 10Hz AV flicker induced more robust pathway modulation compared to 40Hz stimulation. 10Hz AV flicker caused a ∼4-10-fold increase in 15 DEGs that play a role in immune and cell adhesion pathways while 40Hz AV flicker caused a ∼0.5-0.7 Log2-fold change in 19 DEGs associated with metabolic and cellular responses (**Figure 5A-B**, **5D-E, Supplemental Table 8**). Furthermore, 10Hz AV flicker resulted in robust, ∼2-9 Log2-fold downregulation of 47 DEGs involved in structural maturation, metabolic and signaling pathways (**Figure 5A-B, Supplemental Table 2, Supplemental Table 7,** FDR-adjusted p<0.05; see Methods). 40Hz AV flicker modulated 385 DEGs with a ∼2-5 Log2-fold downregulation in similar pathways (**Figure 5D, 5E, Supplemental Table 2, Supplemental Table 8,** FDR-adjusted p<0.05, see Methods). This pattern was also observed in upregulated DEGs. This modulation pattern suggests that 40Hz AV flicker enhances overall astrocytic capacity for broader stress mitigation and synaptic maintenance, while 10Hz AV flicker provides a more targeted restoration of stress-impaired astrocytic functions primarily through modulation in *Fkbp5* signaling pathways as well as metabolic and structural processes.

To corroborate these transcriptional effects of AV flicker on astrocytes, we assessed the effects of stress and AV flicker on astrocyte morphology in the medial prefrontal cortex. Using branching arborization and convex hull analysis to measure the volumetric extension of astrocytes, we observed significant morphological shifts in GFAP-positive astrocytes in response to AV flicker intervention compared to stress alone without stimulation (**Figure 5F**, Males arborization H (3) = 84.70; p<0.0001; Males convex hull H (3) = 43.38; p<0.0001**, Supplemental Figure G,** Females arborization H (3) = 8.856; p=0.0313; Males convex hull H (3) = 3.730; p=0.2922). Notably, 10Hz AV flicker significantly increased astrocyte process length and complexity beyond levels observed in the stressed unstimulated group. This enhanced reactivity suggests a compensatory mechanism to promote resolution of stress-induced alterations, by potentially increasing coverage of synapses to enhance neurotransmitter uptake and metabolic support. This effect was frequency specific as astrocytes morphology after 40Hz AV flicker significantly differed from those after 10Hz AV flicker in stressed males (**Figure 5F**, Dunn’s post hoc males stress 10Hz vs stress 40Hz arborization p<0.0001 and p=0.1510**, Supplemental Figure 9F,** Dunn’s post hoc females stress 10Hz vs stress 40Hz arborization p>0.9999 and p>0.9999). These morphological findings provide additional evidence for frequency-specific modulation of astrocytes in the medial prefrontal cortex in the context of stress.

### Flicker Mitigates Stress-Induced Molecular Profile of Stress Susceptibility in Frontal Cortex Microglia

Microglia are essential for certain stress-induced behavioral changes^63–73^ and play central roles in neuroinflammation and synaptic plasticity under stress conditions^64,67,69,74^. Based on this, we hypothesized that AV flicker that induces behavioral stress resilience would modulate microglial transcriptional responses to stress. To test this hypothesis, we analyzed isolated CD45+ cells which are primarily microglia after animals underwent 10Hz or 40Hz AV flicker and concomitant CUS exposure (**Figure 6, Supplemental Figure 10**). We observed a robust transcriptional response to 10Hz AV flicker stimulation in stressed males with 704 DEGs, compared to 111 DEGs in response to 40Hz AV flicker (**Figure 6, Supplemental Table 3**). AV flicker also altered microglia genes in stressed females with 117 DEGs after 10Hz AV flicker and 6 DEGs after 40Hz AV flicker (**Supplemental Figure 10, Supplemental Table 3**). 10Hz AV flicker enriched genes associated with a broad range of processes in males, including glial development, myelination, and synaptic modulation, indicating that 10Hz AV flicker promotes neuroprotective and homeostatic microglial functions (**Figure 6A, 6B, 6C; Supplemental Table 9;** FDR-adjusted p<0.05; see Methods).

**Figure 6:**
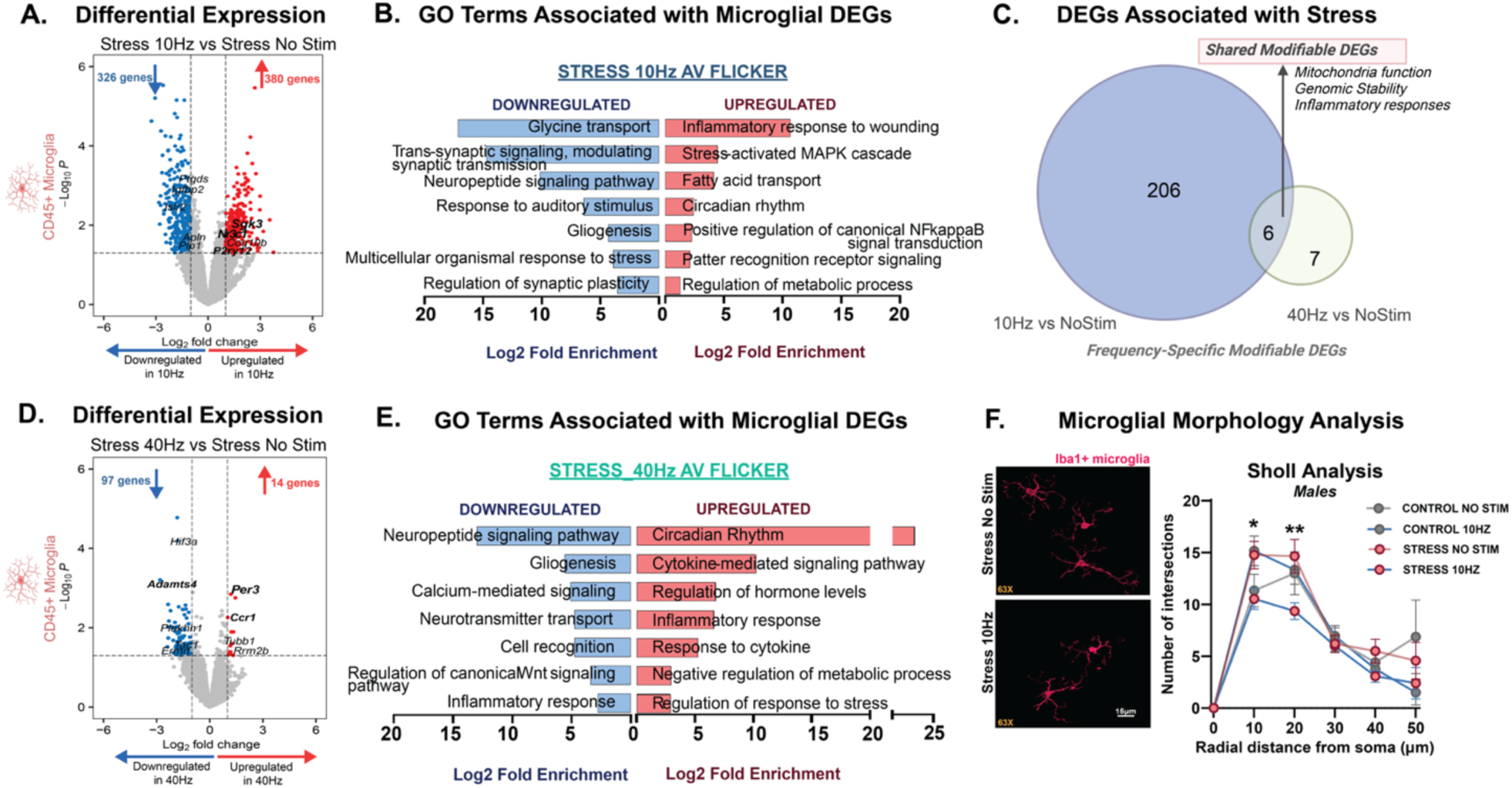
10Hz AV flicker mitigates stress-induced transcriptional changes in frontal cortical microglia. **(A)** Volcano plot showing differential gene expression of microglia in stressed males following 10Hz AV flicker compared to stress no stimulation. Genes involved in the regulation of stress signaling are highlighted in bold. **(B)** Gene ontology (GO) enrichment analysis for genes downregulated (blue) and upregulated (red) after 10Hz AV flicker exposure in stressed males compared to no stimulation stressed males. **(C)** Venn diagram demonstrating unique and shared (pink) modifiable enriched genes associated with stress following 10Hz (blue) AV flicker compared to 40Hz (green) AV flicker. **(D)** and **(E)** As in A and B for 40Hz exposure in stressed males compared to no stimulation stressed males. **(F)** Example 63x image of Iba1+ microglia (magenta) after stress (top left) or stress and 10Hz flicker (bottom left), scale bar 15µm. Cortical microglia branching (assessed via Sholl analysis) in male mice that underwent no stress and no stimulation (grey dot, grey line), no stress and 10Hz AV flicker (grey dot; blue line), stress and no flicker (red dot; red line), or stress and 10Hz flicker (red dot; blue line) conditions (F (5, 489) = 86.96; ***p<0.0001). Error bars show mean ± SEM. N = 3 pooled samples from 6 animals per group for transcriptome analysis and n = 25-31 cells from 3-4 animals per group for morphology analysis. Morphological analyses were conducted at the group level, with data aggregated across cells within a group. Transcriptional profile at gene expression level without FDR adjustment p<0.05, Wald test. GO terms pathway analysis FDR-adjusted p < 0.05. Two-way ANOVA test performed for microglia morphology analysis followed by Bonferroni’s multiple comparisons test. See **Supplemental Table 11** for statistical details.

Consistent with the observed efficacy of 10Hz AV flicker in counteracting stress-induced transcriptional alterations in neurons and astrocytes, we found that 10Hz AV flicker robustly mitigated stress effects in cortical microglia in male mice. Impressively, 30% of DEGs altered by stress were normalized by 10Hz AV flicker intervention, compared to only 12% of DEGs in response to 40Hz flicker (**Supplemental Figure 7A**), showing that 10Hz AV flicker is more effective in mitigating stress-related molecular changes in microglia. GO analysis of downregulated DEGs following 10Hz AV flicker revealed significant terms associated with glycine transport, response to stress and synaptic plasticity (**Figure 6B**, **Supplemental Table 9**, FDR-adjusted p<0.05; see Methods), indicating that 10Hz AV flicker reduces microglial overactivation in these areas. Upregulated pathways at 10Hz predominantly involved inflammatory responses, circadian rhythms, and metabolic processes (**Figure 6B**, **Supplemental Table 9**, FDR-adjusted p<0.05; see Methods), suggesting that 10Hz AV flicker regulated inflammatory state, immune responses, and synaptic maintenance. Interestingly, we found that 10Hz flicker mitigated stress-induced changes in transcription of microglial receptors *P2ry12* and stress associated receptors such as *Nr3c1* and *Tlr4*, as well as microglial *Igfbp2*, a key gene implicated in affective and psychiatric disorders (**Figure 6A, Supplemental Table 3**). These effects were frequency specific and showed little overlap between 10Hz and 40Hz AV flicker (**Figure 6C**). 40Hz AV flicker robustly downregulated signaling pathways involved in inflammatory response and neurotransmission, and upregulated genes related to neurotransmitter regulation and synaptic function (**Figure 6D, 6E**, FDR-adjusted p<0.05; see Methods), indicating that 40Hz AV flicker has a more targeted effect on maintaining synaptic stability and supporting neurotransmission.

To corroborate the observed transcriptional effects of AV flicker, we assessed flicker-mediated modulation of microglial morphology in mPFC. We observed a difference in the branching extension of IBA1^+^ microglia using Sholl analysis following 3D rendering (**Figure 6F,** F (5, 489) = 86.96; p<0.0001). Notably, 10Hz AV flicker in stressed males significantly decreased microglial processes crossing 10-micron (Bonferroni’s post hoc; p=0.0129) and 20-micron (Bonferroni’s post hoc; p=0.0010) distal points from the soma compared to unstimulated stressed males. Decreased branching is consistent with microglia primed for resolution of stress induced cascades. Together, these results show that 10Hz AV flicker attenuates stress-induced changes in microglia transcription and regulates clinically relevant microglial genes. Together, our findings show that 10Hz AV flicker in males robustly affects microglia activity, mitigating stress-induced transcriptional changes and enhancing reparative signals. The effects on microglia in response to 10Hz AV flicker coincided with stress resilience at the behavioral level.

## DISCUSSION

Our findings demonstrate for the first time that AV flicker exposure produces a protective effect in stressed males and females, inducing stress resilience with frequency- and sex-specific effects. Importantly, we found that 10Hz AV flicker in males and 40Hz AV flicker in females were particularly effective frequencies for mitigating stress-induced changes, suggesting a potential for tailored interventions based on sex. Both sexes exhibited reduced anxiety-like behaviors, with males also showing improvements in exploratory behaviors and females demonstrating decreased despair-like behaviors and prevention of stress-induced weight changes at 10Hz and 40Hz, respectively. AV flicker stimulation effectively prevented stress-induced alterations in spine morphology, suggesting a protective effect on synaptic structure. At the molecular level, 10Hz AV flicker in males significantly modulated stress signaling pathways across neurons, astrocytes, and microglia, as indicated by regulation of the norepinephrine receptor gene *Adra2c* in neurons, the glucocorticoid receptor regulator *Fkbp5* in astrocytes, and the glucocorticoid receptor gene *Nr3c1* in microglia **(Supplemental Figure 11)**. Coinciding with this multi-cellular stress signaling regulation, flicker stimulation enhanced pathways associated with synaptic maintenance and plasticity, suggesting a multifaceted neuroprotective effect of AV flicker stimulation in the context of stress. Overall, these findings highlight the promise of AV flicker as a novel strategy for enhancing stress resilience.

We show that optimized frequencies of AV flicker, particularly at 10Hz in males and 40Hz in females, promote protective shifts in glial function. In males, 10Hz AV flicker mitigated stress-induced dysregulation of astrocytic pathways involved in metabolism and synaptic organization, while also reversing microglial activation and enhancing synaptic maintenance pathways. In addition, 10Hz flicker induced more spread of astrocytic processes which may facilitate increased coverage of synapses and blood vessels, potentially enhancing neurotransmitter uptake, metabolic support, and regulation of the blood-brain barrier to restore homeostasis after chronic stress. In females, 40Hz AV flicker enhanced astrocytic pathways related to cellular adhesion and neurotransmitter regulation, with more limited effects on microglia. It remains to be determined whether the observed spine dynamics changes following AV flicker are explained by glia-mediated engulfment. Further investigation, including time-course studies and specific markers of glial engulfment, is needed to elucidate this aspect of the AV flicker-induced effects^6,17,18,26^. Together, the observed glial responses to AV flicker stimulation suggest that astrocytes and microglia serve as key modulators of flicker-mediated neuroprotection, facilitating synaptic stabilization and reducing neuroinflammation under stress conditions.

Notably, 10Hz and 40Hz AV flicker stimulation elicited both overlapping and distinct effects, particularly in neuron-glia communication genes and synaptic remodeling genes. This suggests that while 10Hz and 40Hz AV flicker recruit some common signaling pathways, frequency-specific effects can be used to tailor flicker stimulation to different functions and contexts including pathology severity, sex, or age. We focused the current study on the prefrontal cortex because it is a key stress-sensitive region associated with neuropsychiatric disorders and cognitive decline^6,17,18,26^. Previous preclinical and clinical reports have reported sex differences in neural responses during stress and stress-associated neuropsychiatric disorders in males and females where males showed greater stress responses in prefrontal cortex regions while females had stronger responses in limbic regions, which are also stress-susceptible^75–77^. This aligns with our observations and suggests that the more subtle effect of AV flicker observed in stressed females may be due to regional effects of stress and flicker interaction. Our findings highlight four key pathways involving regulation of stress signaling, inflammatory response pathways, preservation of synaptic integrity and plasticity, and normalization of metabolic and cellular transport processes which motivate future studies to interrogate mechanisms of AV flicker protection. Under chronic stress, excessive excitatory signaling can lead to synaptic damage and neuroinflammation, processes tightly regulated by astrocytes and microglia. AV flicker may enhance glial cells’ capacity to regulate this excitatory-inhibitory balance, thereby preventing the cascade of stress-induced neurotoxicity. Based on our findings, 10Hz AV flicker may preferentially engage astrocytic pathways that support metabolic function and synaptic integrity in males, while targeting microglial processes involved in synaptic pruning and neurotransmitter regulation. In contrast to 10Hz AV flicker, 40Hz AV flicker had broader transcriptional effects but altered fewer stress-modulated genes. Because 10Hz AV flicker in males resulted in the most robust behavioral stress resilience the specific effects of 10Hz AV flicker reflect changes related to stress resilience rather than flicker exposure broadly. Together, our results reveal stress-associated genes and pathways that are modifiable by frequency-optimized non-invasive brain stimulation.

This work motivates several future studies. First, we used immuno-panning together with bulk RNAseq analysis to enable broader quantification of lowly-expressed transcripts and to avoid artifacts associated with tissue dissociation and sorting. Our current approaches may be bolstered by using single nuclear RNAseq, or cell-type specific native state proteomic or transcriptomic approaches. Future research will investigate whether AV flicker reverses established stress-induced deficits if used after chronic stress exposure instead of concomitantly. Moreover, persistence of the observed effects and the potential need for maintenance treatments will be investigated in future studies. It is also worth considering whether acquired resilience following chronic AV flicker exposure mimic innate stress resilience phenotypes. Future studies might benefit from comparing multiple stress models or using stress models in which subsets of subjects show spontaneous innate resilience that could be compared to AV flicker’s effects. Lastly, while our findings provide correlational evidence of protection against stress-induced pathology and behavioral deficits, the causal mechanisms underlying flicker-induced stress resilience remain to be fully elucidated.

In total, our study provides compelling evidence that frequency-specific AV flicker mitigates stress-induced behavioral, synaptic, and molecular changes in both males and females, with distinct effects based on AV flicker frequency. Our findings suggest that AV flicker could serve as a non-invasive, easily implementable intervention for stress-related dysfunction. The frequency-specific and sex-specific effects highlight the possibility of personalized treatment approaches based on individual factors. The different responses in astrocytes and microglia to different flicker frequencies underscores the importance of these cell types in modulating stress resilience and the therapeutic potential of non-invasive neuromodulation techniques. Overall, these findings highlight the promise of AV flicker as a novel strategy for enhancing stress resilience.

## MATERIALS AND METHODS

### Animals and Husbandry

The Georgia Tech Institutional Animal Care and Use Committee (IACUC) approved all animal work performed in this study. For experiments assessing behavior, astrocyte morphology, and transcriptional profiling, adult (2- to 3-month-old) male and female C57BL/6J mice were obtained from the Jackson Laboratory. For experiments assessing neuronal spine morphology and density, adult (2- to 3-month-old) male and female Thy1-EGFP-M mice (https://www.jax.org/strain/007788) were obtained from the Jackson Laboratory^45^. Mice were grouped-housed and maintained in standard conditions with a 12-hour light/dark cycle. Animal housing rooms were equipped with a ventilation system that provides 12 air changes per hour, temperature range of 64-79°F and 30-70% relative humidity. Animals had ad libitum access to food and water except during food or water deprivation stressors. All experiments were conducted during the light cycle at various time points^78^.

### Chronic Unpredictable Stress Paradigm

To induce sustained psychological stress in the experimental mice, animals were subjected to a chronic unpredictable stress (CUS) paradigm was implemented over a standardized period of 21 consecutive days (n=6-9 mice/group). In some studies, the exposure period was extended by an additional 7 days to accommodate behavioral testing. Body weight was tracked longitudinally from the first day of stress exposure until the last day of the paradigm. Each day, the animals were subjected to two distinct mild to moderate stressors, presented at random intervals between 8 AM and 5 PM to maintain the unpredictable nature of the stress induction. For stressors requiring extended exposure, such as overnight manipulations, the stress period was extended from 5 PM to 8 AM the following day. The CUS protocol included a variety of physical and psychological challenges: overnight water deprivation, overnight food deprivation, reversed light/dark cycles over a 24-hour period, overnight exposure to wet bedding, 1-hour periods of restraint stress, 45-minute intermittent cage shaking, 10% peppermint oil exposure, and periods of radio static noise. As the final stressor, all stress-exposed mice received 1-hour restraint stress in the morning (**Supplementary Figure 1**). Tissue samples were collected 60-90 minutes post-restraint, concluding the experimental stress protocol. At the end of the 4-week CUS exposure, behavioral and physiological assessments were conducted, including measurements of body weight, anhedonia (via the sucrose consumption test), despair (via the forced swim test), anxiety-like behaviors (via the elevated plus maze and open field test), as well as general exploration, locomotion, and recognition memory (via the novel object recognition test). To quantify the overall impact of CUS, we employed an adapted version of the composite behavioral and physiological score developed by Johnson et. al (2021). This score incorporated behavior tests that map on to the research domain criteria (RDoC)^43^ which revealed a robust increase in stress-induced maladaptive behaviors across both sexes. This comprehensive score integrated averaged z-scored outcome measures from all the aforementioned behavioral and physiological tests (assays and scoring described below under Behavior Screening). The resulting susceptible-resilience score was normalized to stress-naïve controls with sex-specific directionality indicating an increased resilience to stress and negative values signifying heightened stress susceptible. The composite susceptible-resilience stress score for females included an additional social interaction assessment using the three-chamber social interaction test due to the higher variability in standard stress assays observed in females. This additional assay approach allowed us to better capture female stress in known behaviors affected in females.

### Audiovisual flicker Stimulation

Animals were exposed to audiovisual (AV) flicker stimulation at one stimulation frequency (40Hz, 20Hz, or 10Hz) daily for 1 hour per day over the course of the CUS paradigm. Animals were randomly assigned to AV flicker exposure groups, with experimenters blinded to the conditions during analysis. Flicker exposure occurred on the same days as CUS but not at the same time. The AV flicker stimulation was administered within a standardized window between 8 AM and 5 PM, either before or after the CUS stressor presentation. For the AV flicker exposure, mice were transferred daily to an empty cage with three sides blocked by matted black walls and one clear side facing a strip of LEDs. The animals were subjected to one of four conditions: LED lights flashing at 40Hz frequency (12.5-ms light on/off cycles), 20Hz frequency (25-ms light on/off cycles), 10Hz frequency (50-ms light on/off cycles), or no stimulation (0 Hz, constant darkness as control). Each exposure session lasted 1 hour, with LED intensity set at 100-400 lux at the animal’s head level and coupled with frequency specific sound presentation set at 65 decibels on average. These specific frequencies were chosen based on previous research indicating their potential effects on neural oscillations, neuroimmune function, and behavioral improvement^11,12,39,40,79–81^. Daily AV flicker stimulation was maintained throughout the entire 21-30-day experimental period, coinciding with the duration of the CUS protocol. By integrating this AV flicker stimulation protocol with the CUS paradigm, the study aimed to investigate potential protective or modulatory effects of specific light frequencies on stress-induced neurobiological and behavioral changes.^81^

### Behavior Screening

#### Composite susceptibility score

We employed an integrative z-scoring method to combine the physiological behavioral readouts into a single score for each behavioral dimension per animal^43^. The z-score for each individual test was calculated based on the standard deviation to experimental group and mean of the control group, separated by sex.

The composite behavioral and physiological stress score was calculated as follows^43^:

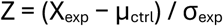

Where: X_exp_ is the individual value from the experimental group, μ_ctrl_ is the mean (average) of the control group, σ_exp_ is the standard deviation of the experimental group. This formula standardizes how far an individual experimental value deviates from the control group mean, scaled by the variability within the experimental group.

#### Sucrose Consumption Test

This test is used as a consistent measure of anhedonia-like behavior^82^. Mice were habituated to 1% sucrose solution for 48hrs with ad libitum access to sucrose solution from Day 19-21 of chronic unpredictable stress, with no access to other fluids. On Day 21, mice were tested for overnight sucrose consumption with 1% sucrose solution, following 12hrs of fluid restriction. Overnight sucrose consumption was measured for two consecutive days (Day 21-23). On Day 24, mice were fluid restricted for 12hrs and then tested for overnight water consumption. Following the sucrose consumption test, daily water intake was tracked until the end of the paradigm.

#### Forced Swim Test

This test is used as a measure of despair-like behavior and antidepressant effects of drugs and treatments in rodents^83^. On Day 25 of chronic unpredictable stress, each animal was placed in a clear swim cylinder and subjected to one 10-minute session of swimming in water while being recorded from above with a ceiling mounted camera. Immobility time was hand-scored by a blinded observer during a 4-minute testing period, with the first 2 minutes of habituation excluded from analysis. Immobility was defined as the least amount of movement possible to stay afloat.

#### Elevated Plus Maze

This test is used as a consistent measure of exploration and anxiety-like behavior. On Day 26 of chronic unpredictable stress, each animal was placed at the center cross point of the elevated plus shaped apparatus (Noldus), with two open arms and two enclosed arms in a dimly lit room. Mice were allowed to explore freely explore all arms for 6 minutes. Maze position, time, and number of entries in the open arms are recorded from above with a mounted camera and quantified using Noldus Ethovision software (Ethovision 15).

#### Open Field Test

This test is used as a consistent measure of exploration and anxiety-like behavior^84^. On Day 27 of chronic unpredictable stress, mice were placed in an open novel arena (Noldus) and allowed to explore for 15 minutes. Maze position, time, and number of entries in the center zone were recorded from above with a mounted camera and quantified using Noldus Ethovision software (Ethovision 15).

#### Locomotor Activity

Ambulatory changes that may influence maze exploration were screened for in the open field maze and quantified as locomotor activity^82–84^. This screen was conducted during maze exploration in the empty open field prior to novel object recognition. During the assay mice were allowed to freely roam in the open field for 20 minutes. Distance travelled in meters was quantified using Noldus Ethovision software (Ethovision 15).

#### Additional behavioral tests

Male mice were assessed for object recognition using the novel object recognition test, while female mice were tested for social avoidance in the social interaction test. In the microglia depletion study, which utilized the colony-stimulating factor 1 receptor (CSF1R) inhibitor PLX3397, male mice were also evaluated using the novelty-suppressed feeding test Specifically, testing in PLX3397- and vehicle-exposed mice was started 1 days earlier (testing was done on CUS days 20-29) enabling male mice to be assessed in the novelty-suppressed feeding test.

#### Novel Object Recognition Test

This test is used as a measure of object recognition memory^82^. On Day 28 of CUS, following maze habituation (see above), male mice were placed in an open novel arena (Noldus) and exposed to two identical objects positioned on opposite sides of the arena. Mice were allowed to explore for 15 minutes. The objects were placed away from the walls, ensuring ample space for the mice to move around them freely. After a 20-hour inter-trial interval on Day 29 of CUS, mice were reintroduced to the arena, where one of the original objects (familiar object) remained in the same location, while a new object (novel object) replaced the second original object. The placement of the novel and familiar object was counterbalanced across trials to avoid spatial bias. The position of the arena and time spent interacting with each object were recorded from above using a mounted camera and analyzed with Noldus EthoVision 15 software. A discrimination index was calculated to quantify preference for novel interaction relative to familiar object exploration.

#### Social Interaction Test

This test assesses social interest and social avoidance^85^. On Day 28 of CUS, female mice were placed in a three-chamber arena (Noldus), which consists of three equal-sized compartments with open doorways allowing free movement between chambers. Mice began in the center chamber and were given 10 minutes to explore both the social chamber, containing an unfamiliar young adult female mouse enclosed in a ventilated wire cage, and the object chamber, containing an inert novel object in an identical cage. The placement of the social and object chambers was counterbalanced across trials to avoid spatial bias. To ensure unrestricted movement, the cages were positioned away from the walls. Exploration behavior (including time spent interacting with the unfamiliar mouse versus the inert object) was recorded from above using a mounted camera and analyzed with Noldus EthoVision 15 software. A discrimination index was calculated to quantify preference for social interaction relative to object exploration.

#### Novelty-Suppressed Feeding

This test assesses anxiety-like behavior and the motivation to eat in a novel environment^84^. The NSF test began after an overnight 16-hour food deprivation period. On Day 27 of CUS, male mice were placed in a novel open arena under dimly lit conditions, with a single food pellet positioned at the center. A blind experimenter manually recorded the latency to approach and consume the pellet. All trials were also recorded from above using a mounted camera and analyzed with Noldus EthoVision 15 software. After completing the novel environment test, mice were immediately transferred back to their home cage, where food pellets were placed in a dish at the center. The latency to feed in the home cage was recorded to control for potential confounds related to appetite drive, ensuring that feeding latency in the novel arena reflects anxiety-related avoidance rather than differences in hunger levels.

### Microglia Depletion

Pexidartinib (PLX3397, MedChemExpress, **Supplemental Table 10**), a colony-stimulating factor 1 receptor (CSF1R) inhibitor, was used to deplete microglia in the central nervous system^12,86,87^. PLX3397 was incorporated into an open standard diet containing 15 kcal% fat, supplied by Research Diets Inc., at a dose of 290 mg/kg. A control group was maintained on a calorie-matched open standard diet, also with 15 kcal% fat content. Mice were fed either depletion or control diet during the pre-stress period (3 days prior) and the respective diets were continued throughout the stress paradigm and the behavioral screening phase (n=6/group).^12^

### Histology

All brain tissue was collected between the hours of 9 AM-1 PM. Mice were deeply anesthetized (5% isoflurane) and subjected to transcardiac perfusion with 4% paraformaldehyde (PFA) to rapidly fix the brain tissue in situ. Whole paraformaldehyde (PFA)-fixed brains were extracted. Following extraction, the brains underwent an additional 48-hour post-fixation period in cold 4% PFA to ensure thorough tissue fixation. After fixation, each hemisphere was rinsed and placed in 30% sucrose for 3 days for cryoprotection and then frozen. Coronal sections were obtained at 30 μm and stored in glycerol-based cryoprotectant solution. The sections underwent a series of rinses in 1X PBS, followed by blocking in a buffer containing 0.3% Triton X-100, 7.5% normal goat/donkey serum, and 1X PBS. Primary antibodies (anti–GFAP and anti–Iba-1, **Supplemental Table 10**) were applied overnight at 4°C, followed by secondary antibody incubation (goat anti-chicken 555 and goat anti-mouse 488, **Supplemental Table 10**) overnight at 4°C in blocking solution. Nuclei were stained with 4′,6-diamidino-2-phenylindole (DAPI) for 1 minute, and sections were rinsed with 1XPBS before being mounted using VECTASHIELD. Sections were imaged using a Nikon Crest spinning disk confocal microscope with a 20X objective for glial imaging and reconstruction for astrocyte [Image size: x:512 y:512 z:25-30 microns; Voxel size: 0.6252×0.6252×1 micron^3], and Inverted Abberior Super-Resolution using oil 63X objective for neuronal spine morphology assessment [Image size: ×:750 y:750 z:44-50 microns; Voxel size: 0.1667×0.1667×0.5000 micron^3]. Imaging focused specifically on the medial prefrontal cortex (mPFC), a region crucial for higher-order cognitive functions and often implicated in stress-related disorders. Within the mPFC, the imaging spanned layers I through V, capturing the full cortical depth and allowing for comprehensive analysis of neuronal architecture across different functional layers. Each z-stacked image generated from stitched confocal images provided a high-resolution snapshot of this region.

### Cellular Morphology

Bitplane Imaris 9.7.1 (Oxford Instruments, Concord, MA) was employed for 3-dimensional (3D) rendering of Thy1+ neuronal spine, GFAP+ astrocyte, and Iba1+ microglia morphology, with the experimenter blinded to experimental groups. Surface reconstruction parameters were optimized across samples using Imaris integrated machine learning (ML) approaches and visually examined while blinded and before hypothesis testing. Each mouse was sampled a minimum of two times. Morphological analyses were conducted at the group level, with data aggregated across cells within a group rather than computing an average value per subject.

*<u>Cortical pyramidal neuron analysis</u>*, 3D rendering of segments in layer II/III of the mPFC was performed using Imaris software. This process utilized high-resolution image stacks of Thy1-EGFP labeled neurons acquired through super-resolution microscopy. Following detailed 3D renderings of neuronal structures, spine morphology was categorized into four types: long thin, filopodia, stubby, and mushroom spines. This classification was based on spine length, head diameter, and neck width. Spine density was quantified as the number of spines per micron. One dendrite was analyzed per cell, and minimum 5 dendrites per group were analyzed by a blinded rater.^23^*<u>Astrocyte and</u> <u>microglia morphology analysis</u>,* astrocytes and microglia whose processes were completely within the image bounds were automatically reconstructed using standardized thresholds for signal cluster diameter and surface detail level. Post-reconstruction, surfaces were masked to filter out any remnants not part of a reconstructed cell. Surfaces were then transformed into processes based on standardized thresholds including diameter, gaps between surfaces, and surface detail, followed by quantification of process properties. Convex hull and arborization were measured for volumetric cell extension and total branching length respectively and served as a proxy for cell reactivity and surveillance state.^12^

### Immunopanning

Immunopanning was employed to isolate enriched populations of microglia (CD45+), astrocytes (HepaCam+), and neurons (Thy1+) from frontal cortical tissue encompassing the mPFC, following methods previously published ^55–57^. This approach was chosen to ensure sensitivity and abundance of transcripts while enabling detailed investigations into cell-type-specific transcriptional changes. Antibody-coated plates were prepared at least one day prior to immunopanning. Panning plates were prepared by coating petri-dishes with a secondary antibody the night before, and adding the cell marker specific primary antibodies an hour before immuno-panning. On the day of the procedure, mouse brains were extracted, cortices microdissected, and meninges removed. The tissue was then enzymatically and mechanically dissociated to create a single-cell suspension. Transcription and translation inhibitors including Actinomycin D (5 ug/mL), Triptolide (10uM), and Anisomycin (27.1 ug/mL) were also present in the dissociation to prevent ex-vivo expression artifacts^88^. This suspension was systematically passed over a series of antibody-coated plates, each prepared with specialized antibodies targeting distinct cellular markers (**Supplemental Table 10**). Because the cell suspension covered the antibody-coated plates, targeted cells bound specifically to their corresponding antibodies, while non-specific cells were washed away. This process highly enriched cell populations, often exceeding 95% and with minimal compromise to cellular viability or functional integrity. The recovered cells were then gently detached, pooled into 2 subjects per sample to ensure sufficient sensitivity to detect lowly abundant immune genes, and centrifuged to form cell pellets. Cell pellets were collected for subsequent RNA extraction and RNAseq analyses.^55–57^

### Transcriptomic Analysis

RNA was extracted from immunopanned enriched cell pellets using the Qiagen RNeasy kit (Qiagen; Hilden, Germany, **Supplemental Table 10**) following manufacturer protocols. RNA transcriptomes were quantified from 3 samples (6 subjects with 2 subjects pooled per sample) per sex per condition. Transcriptomic analysis was performed separately for each cell type—neurons, astrocytes, and microglia. Genes with fewer than 50 raw counts in at least 75% of samples were excluded from analysis. Non-coding genes were also removed. Gene counts were normalized using the DESeq2 package in R. Differential expression analysis was conducted using DESeq2to identify DEGs, "*Stress*" was included as a variable in the design matrix when comparing the stress versus control conditions. Similarly, "*Frequency*" was used as a variable in the design matrix to compare different stimulation frequencies against the no-stim condition in stressed-induced subjects. Significant level (p value, Wald test) and Fold change (FC) were computed using *DESeq function.* |Log2FC| ≥1 and p<0.05 were considered as significant DEGs and displayed as volcano plot using *EnhancedVolcano* function in R^89^. Gene ontology analysis on DEGs within each cell type was conducted using PANTHER overrepresentation test with PANTHER 18.0 through the Gene Ontology resource (https://geneontology.org/) against all genes within the dataset. The *Mus Musculus* GO biological processes complete annotation set was used with Fisher’s exact test with FDR adjustment for AV flicker compared to stress no stimulation, or unadjusted for stress no stimulation compared to no stress no stimulation control groups to compute significance of the gene sets for set of DEGs. PANTHER overrepresentation tests against the mouse reference genome mm10 GO biological processes complete annotation set. Gene set variation analysis (GSVA) was conducted to assess functional changes within specific cell types using custom gene sets for neurons using a modified list from previously published sets. GSVA is an unsupervised enrichment algorithm which identifies variations of pathway activity by defining enrichment score for gene sets which each contain a set of genes that share same cellular function. GSVA was performed using R and the GSVA package available on Bioconductor to identify enrichment within neuronal transcriptomes^59,90^.

Modifiable markers were defined as genes upregulated by stress and downregulated by flicker, or vice versa. The percentage of modifiable markers that were altered by flicker stimulation was calculated as: (frequency-specific modifiable DEGs/frequency-specific DEGs)x100 for each frequency.

### Statistical Analysis

For behavior and astrocyte morphology analyses, normally distributed data were analyzed using one-way ANOVA, while non-normally distributed data were analyzed using the Kruskal-Wallis test, with an α value of 0.05 followed by Bonferroni correction for multiple comparisons or Dunn’s multiple comparisons test (respectively) to assess significant differences between groups. For dendritic spine analysis, two-way ANOVA with an α value of 0.05 was used to determine significant interaction of spine class and experimental groups for morphology analysis or interaction of stress and flicker for spine density analysis, followed by uncorrected Fisher’s LSD for multiple comparisons to assess significant differences between groups. For microglia morphology analysis, two-way ANOVA with an α value of 0.05 was used to determine significant interaction of stress and flicker, followed by Bonferroni correction for multiple comparisons to assess significant differences between groups. Compiled stress susceptible score and individual behavior tests were analyzed with independent ANOVA analyses using GraphPad Prism 8 (GraphPad Software, La Jolla, CA) to determine statistical significance between groups. Bonferroni correction for multiple comparisons was applied to determine differences between specific groups^42,44,46^, and uncorrected Fisher’s LSD was applied for multiple comparisons in the spine density dataset. Statistical details for each analysis are described in **Supplemental Table 11**.

## Data and code availability

Morphology data will be uploaded onto Neuromorph.org upon publication. Transcriptomic data is available on NCBI GEO via NCBI Gene Expression Omnibus accession number GSE298961. Custom code used in this study is available on GitHub via https://github.com/singerlabgt/FlickerStressMitigation.git.

## Acknowledgements

We thank Srikant Rangaraju and the Rangaraju lab for useful discussions about this study and the Singer lab for valuable comments on the manuscript. We thank the Emory University Integrated Cellular Imaging Core and Georgia Tech Optical Microscopy Core for access and assistance with confocal imaging. We thank Jacob Tharayil, Jacob Kraus, Ophelia Winslett, and Sara Pennebaker for technical assistance. C.R., N.K., and T.G. was supported by Georgia Institute of Technology President’s Undergraduate Research Awards. A.P. was supported by Atlanta Veterans Affairs RRCD CDA2 E4544-W and Emory Goizueta Alzheimer’s Disease Research Center (ADRC) funded via the National Institute of Aging grant. A.C.S. was supported by the BrightFocus Foundation Grant A2022048S, Emory Goizueta Alzheimer’s Disease Research Center (ADRC) funded via the National Institute of Aging grant P30 AG066511, the Packard Foundation, the National Institutes of Health (NIH)-National Institute of Neurological Disorders and Stroke Grant R01 NS109226 and 2RF1NS109226, the NIH National Institute of Aging Grant RF1AG078736, McCamish Foundation, Friends and Alumni of Georgia Tech, and the Lane Family. T.C.F was supported by the BrightFocus Foundation Grant A2022048S, Emory Goizueta Alzheimer’s Disease Research Center (ADRC) funded via the National Institute of Aging grant P30 AG066511, and the Alzheimer’s Association Research Fellowship AARFD-21-853104. SAS was supported by NIMH R01 MH125956. L.B.W and S.B. were funded in part by the George W. Woodruff School of Mechanical Engineering Woodruff Fellowship.

## Author contributions

TCF: Study conceptualization, funding acquisition, data collection, data analysis, manuscript writing and editing. SB: Data analysis, manuscript writing and editing. ATK: Methodology development, data collection. MG: Data collection. CR: Data collection, data analysis. TG: Methodology development, data collection. AP: Data analysis, manuscript writing and editing. HZ: Data collection, data analysis. NK: Data collection, data analysis. SAS: Study conceptualization, manuscript editing. LW: Study conceptualization, manuscript writing and editing. ACS: Study conceptualization, funding acquisition, data analysis, manuscript writing and editing, supervision.

