## Supplemental Figures for "Sensory neurostimulation promotes stress resilience with frequency-specificity"

for

**Contents**

13 Supplemental Figures: S1-13 (in this file)
11 Supplemental Tables: Table 1-11 (captions in this file, tables are each separate files).
Tables include Key Resources Table (Table 10) and Statistical Details (Table S11).

**SUPPLEMENTARY FIGURES**

Supplemental Figure 1

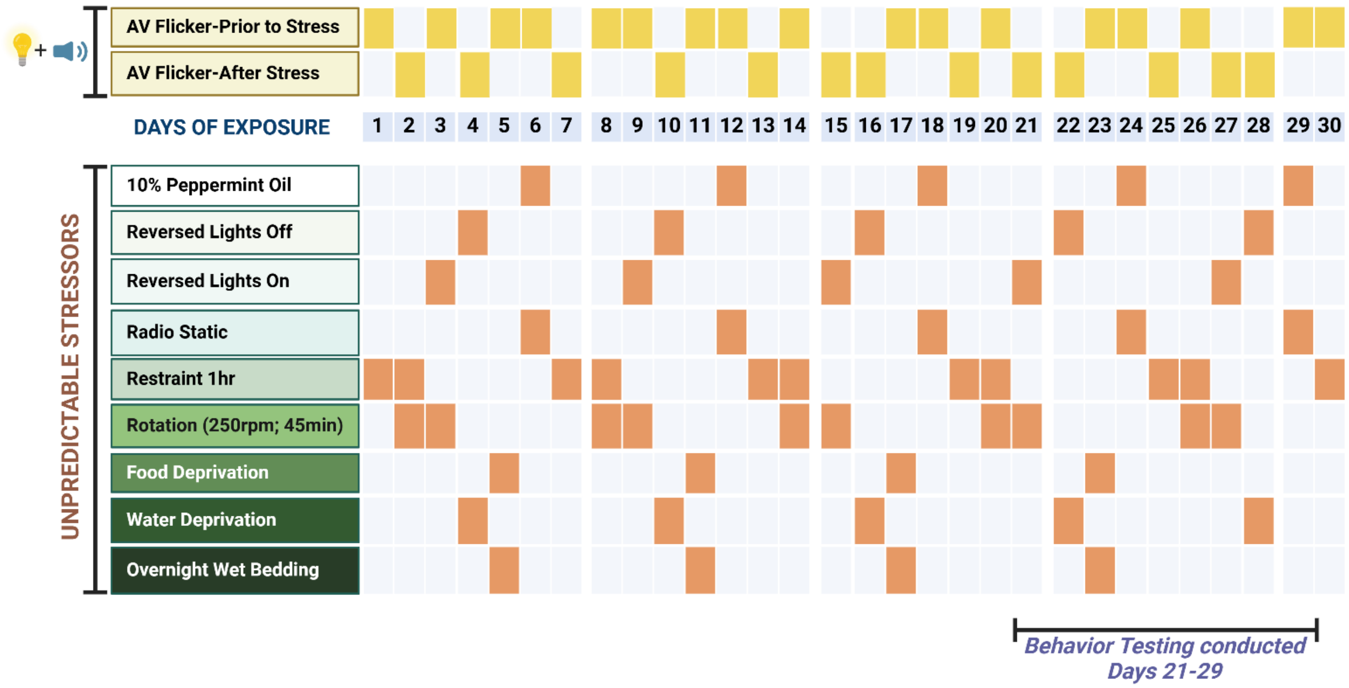


**Supplemental Figure 1. Example Schedule for Concomitant Stress and Audiovisual (AV) Flicker Paradigm.** This table shows an example daily schedule for concurrent exposure to unpredictable stress (stressors listed on the left) and AV flicker (1 hour daily at 10Hz, 20Hz, or 40Hz), with the stressor used each day marked in orange in the bottom rows. The AV flicker was presented in the morning, either before or after stress exposure (marked in yellow in the top rows), in both male and female mice. Created in BioRender. Franklin, T. (2025) https://BioRender.com/v13k485

Supplemental Figure 2

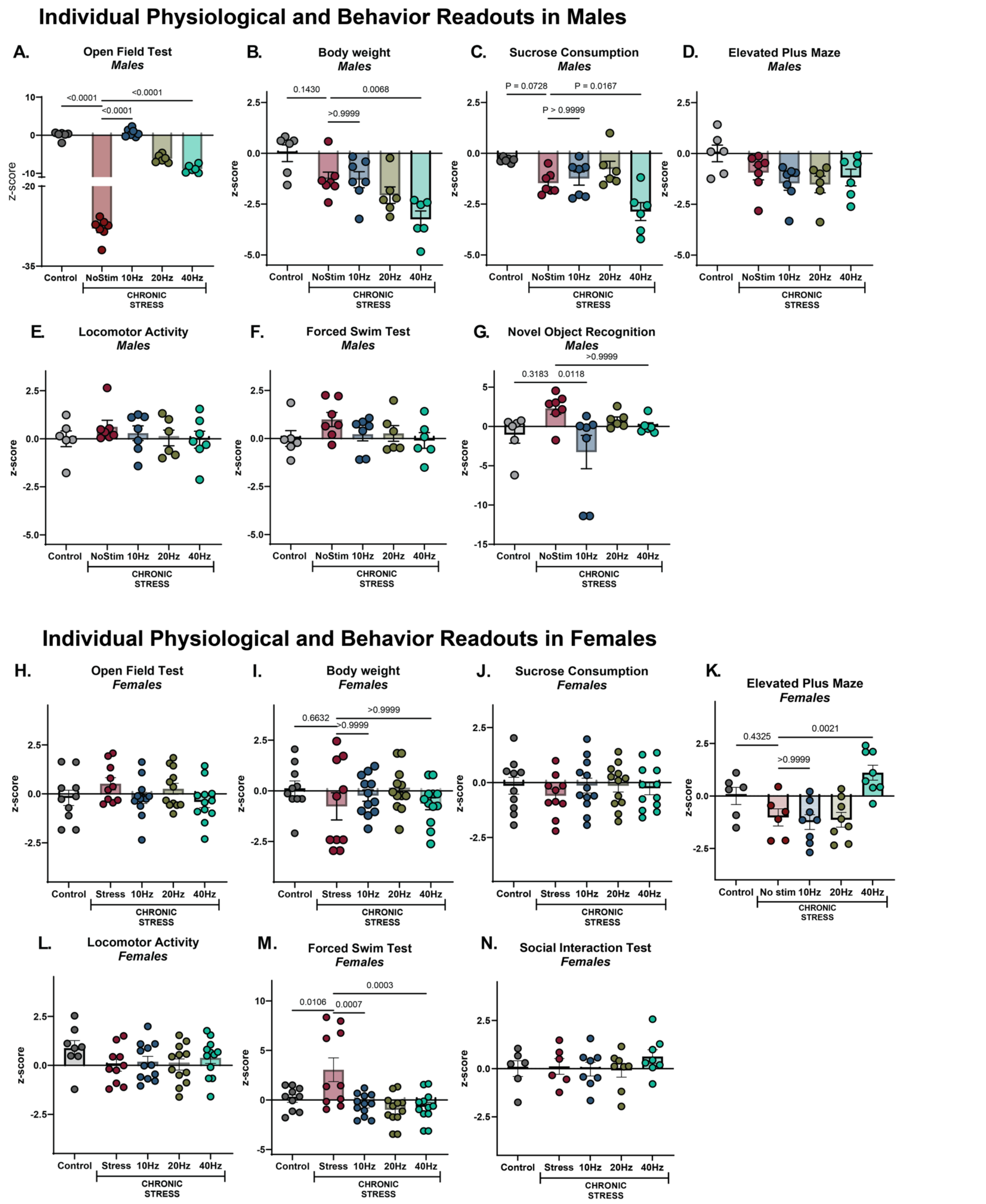


**Supplemental Figure 2: Chronic AV Flicker boosts resilience to stress in male and female mice.** (**A-G**) Stress induced behavioral shifts in male mice that underwent 10Hz (blue), 20Hz (brown), 40Hz (green), stress alone (red), or no stress no stimulation control (grey) conditions in **(A)** open field test ($F_{(4, 27)} = 553.5$; $p<0.0001$), **(B)** body weight ($F_{(4, 27)} = 8.691$; $p<0.0001$), **(C),** sucrose consumption ($F_{(4, 27)} = 9.076$; $p<0.0001$) **(D)** elevated plus maze ($F_{(4, 27)} = 2.503$; $p=0.0659$), **(E)** locomotor activity ($F_{(4, 28)} = 0.5030$; $p=0.7338$), **(F)** forced swim test ($F_{(4, 27)} = 1.276$; $p=0.3038$), and **(G)** novel object recognition ($F_{(4, 27)} = 3.124$; $p=0.031$). 10Hz AV flicker in males reverse or mitigated stress-induced changes in the open field, forced swim, and novel object recognition test although the effects were only sometimes significant when considering each behavior alone. In some assays like body weight and sucrose consumption, 10Hz AV flicker shifted mitigated stress effects in some animals but not others, highlighting the individual responses to stress and the need to consider composite stress scores. (**H-M)** As in A-F for females. (**N**) Social interaction in females ($F_{(4, 31)} = 0.6652$; $p=0.6209$). In females the effects of stress alone compared to no stress were less pronounced than in males, perhaps because of higher variability between individuals. 40Hz flicker in females mitigates the effects of stress on the open field ($F_{(4, 31)} = 1.894$; $p=0.1364$), sucrose consumption ($F_{(4, 31)} = 1.331$; $p=0.2804$), elevated plus maze ($F_{(4, 31)} = 7.861$; $p=0.0002$), locomotor activity ($F_{(4, 31)} = 1.139$; $p=0.3563$), and forced swim test ($F_{(4, 31)} = 2.937$; $p=0.0361$) although the effects were only sometimes significant when considering each behavior alone. Body weight changes were observed in stressed female compared to no stress control, but were not mitigated by 40Hz ($F_{(4, 31)} = 8.011$; $P<0.0001$). Error bars show mean ± SEM; n = 6-8 animals per group. One-way analysis of variance performed for compiled scores followed by Bonferroni correction for multiple comparisons.

Supplemental Figure 3

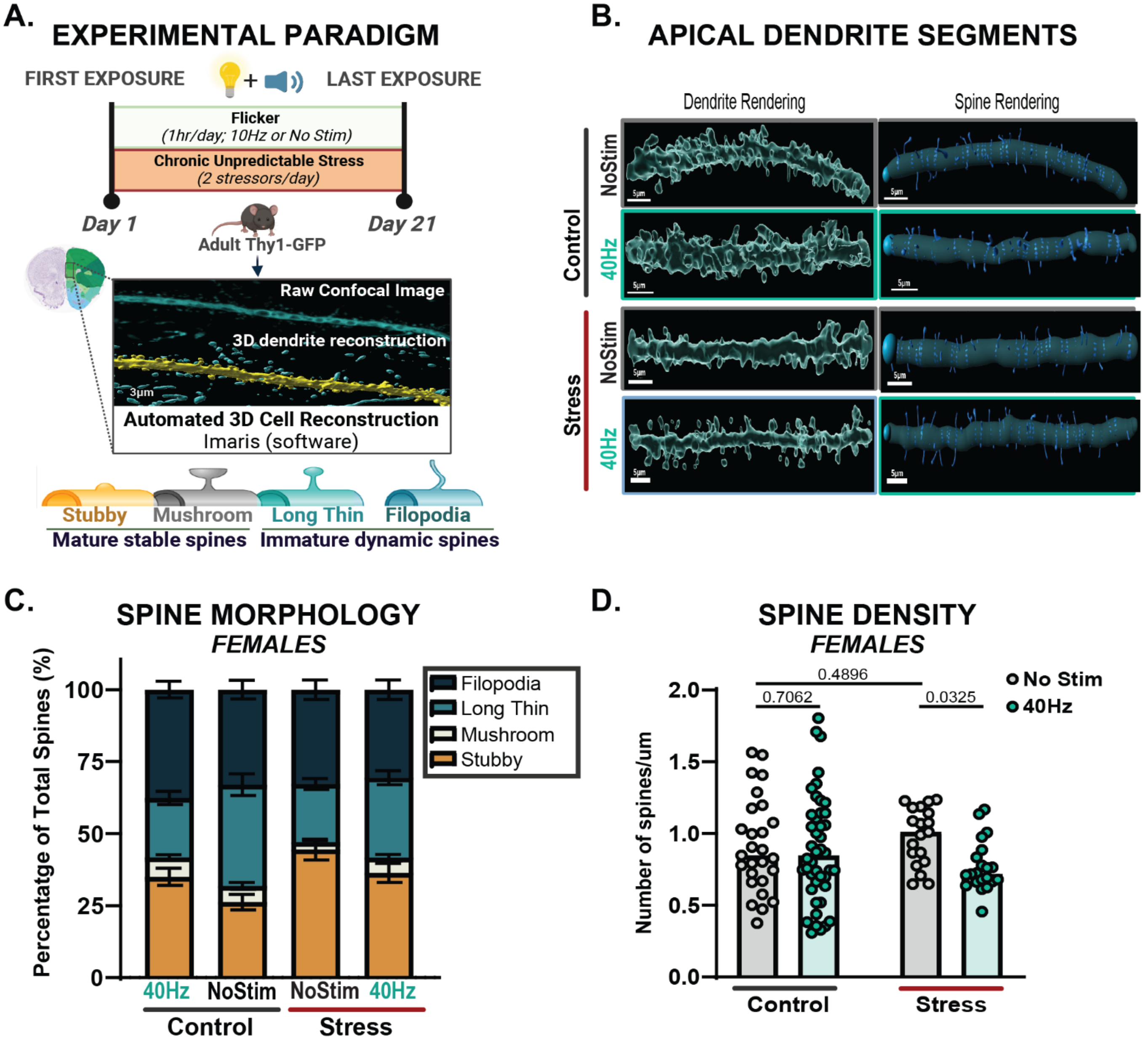


**Supplemental Figure 3: Chronic AV Flicker maintains ratio of dynamic and stable cortical dendritic spines of stressed female mice. (A)** Top: Experimental paradigm. Bottom: Super-resolution confocal images and 3D morphology reconstruction of apical dendritic spines in Imaris imaging software software identified mature stable spines including stubby (orange) and mushroom (grey), and immature dynamic spines including long thin (light turquoise) and filopodia (dark turquoise). **(B)** Representative images of targeted apical dendrite segments from layer II/III of medial prefrontal cortex. **(C)** Spine morphology of each type in female mice following either no flicker ("NoStim") or 40Hz AV flicker ("40Hz") in no stressed ("Control") and CUS ("Stress") conditions ($F(9, 428) = 4.814$, $p<0.0001$). 40Hz AV flicker normalizes immature to mature spine ratio in stressed females exposed compared to stress no stimulation group ($F(1, 107) = 5.010$; $p = 0.0273$) driven by the maintenance of long thin spines ($F(1, 107) = 13.88$; $p = 0.0003$). **(D)** Bar graph showing spine density in female mice following either no flicker (no stim, grey; or 40Hz AV flicker (40Hz, green) in healthy and stress conditions ($F(1, 107) = 3.785$, $p=0.0543$). Each dot is a dendrite segment. Error bars show mean ± SEM; n = 19-44 dendrite segments from 4-6 animals per group. Morphological analyses were conducted at the group level, with data aggregated across dendrite segments rather than averaged per subject. Two-way and one-way analysis of variance performed for compiled scores followed by Bonferroni correction for multiple comparisons for spine morphology and spine density, respectively.

Supplemental Figure 4

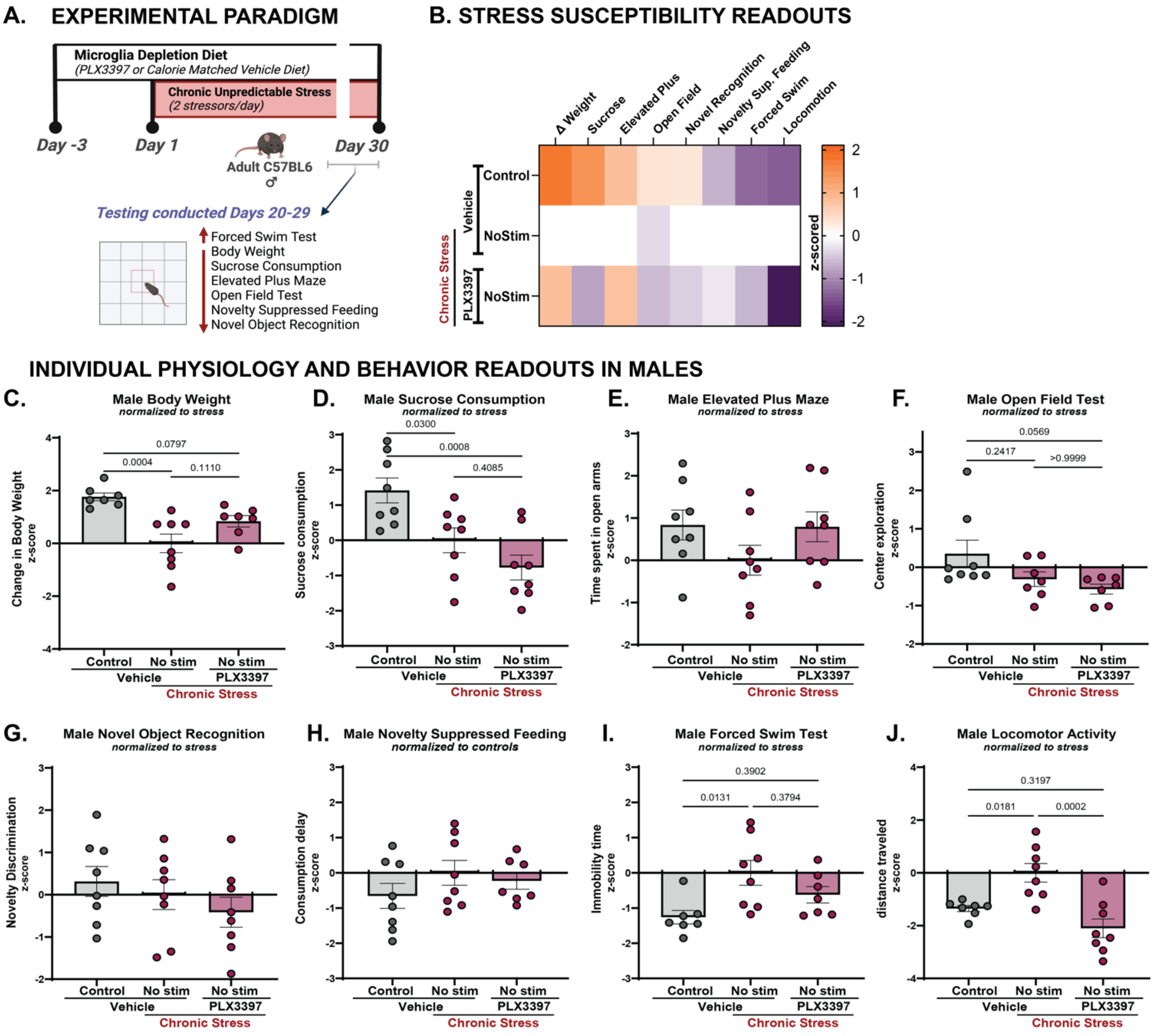


**Supplemental Figure 4: Development of stress-induced susceptible phenotype requires microglia. (A)** Male mice were exposed to microglia depletion diet (PLX3397) or calorie matched vehicle diet concomitantly with daily stress exposure. Control animals (grey) underwent no depletion and no stress exposure. Below: mice underwent a battery of behavioral assays in the last week of stress exposure. **(B)** Matrix of microglia-depleted subjects and no drug controls, showing changes in behavior relative to the baseline stress no stimulation, no drug group. Increases in behavior are indicated in orange, while decreases are shown in purple, reflecting shifts from the baseline score. **(C-J)** Box plots of performance in anxiety-related behaviors and depressive-like behaviors in stressed (red) male mice with ("PLX3397") or without ("Vehicle") microglia depletion colony stimulation factor receptor 1 inhibitor drug PLX3397 compared to no stress vehicle control (grey) **(C)** Change in body weight ($F(2, 19) = 11.17$, $p=0.0006$) **(D)** sucrose consumption ($F(2, 21) = 9.867$, $p=0.0010$) **(E)** elevated plus maze ($F(2, 21) = 1.769$, $p=0.1951$) **(F)** open field test ($F(2, 19) = 3.557$, $p=0.0487$) **(G)** novel object recognition ($F(2, 21) = 1.070$, $p=0.3610$) **(H)** novelty suppressed feeding ($F(2, 20) = 1.084$, $p=0.3574$) **(I)** forced swim test ($F(2, 19) = 5.230$, $p=0.0155$) **(J)** locomotion activity ($F(2, 20) = 12.45$, $p=0.0003$). Microglia depletion with PLX3397 attenuated stress responses in body weight, forced swim test, and elevated plus maze performance, while depletion exaggerated or did not disrupt stress-induced changes in the open field test center entries and sucrose consumption. The results are expressed as the mean ± SEM; n = 7-8 animals per group. One-way analysis of variance performed for compiled scores followed by Bonferroni correction for multiple comparisons.

Supplemental Figure 5

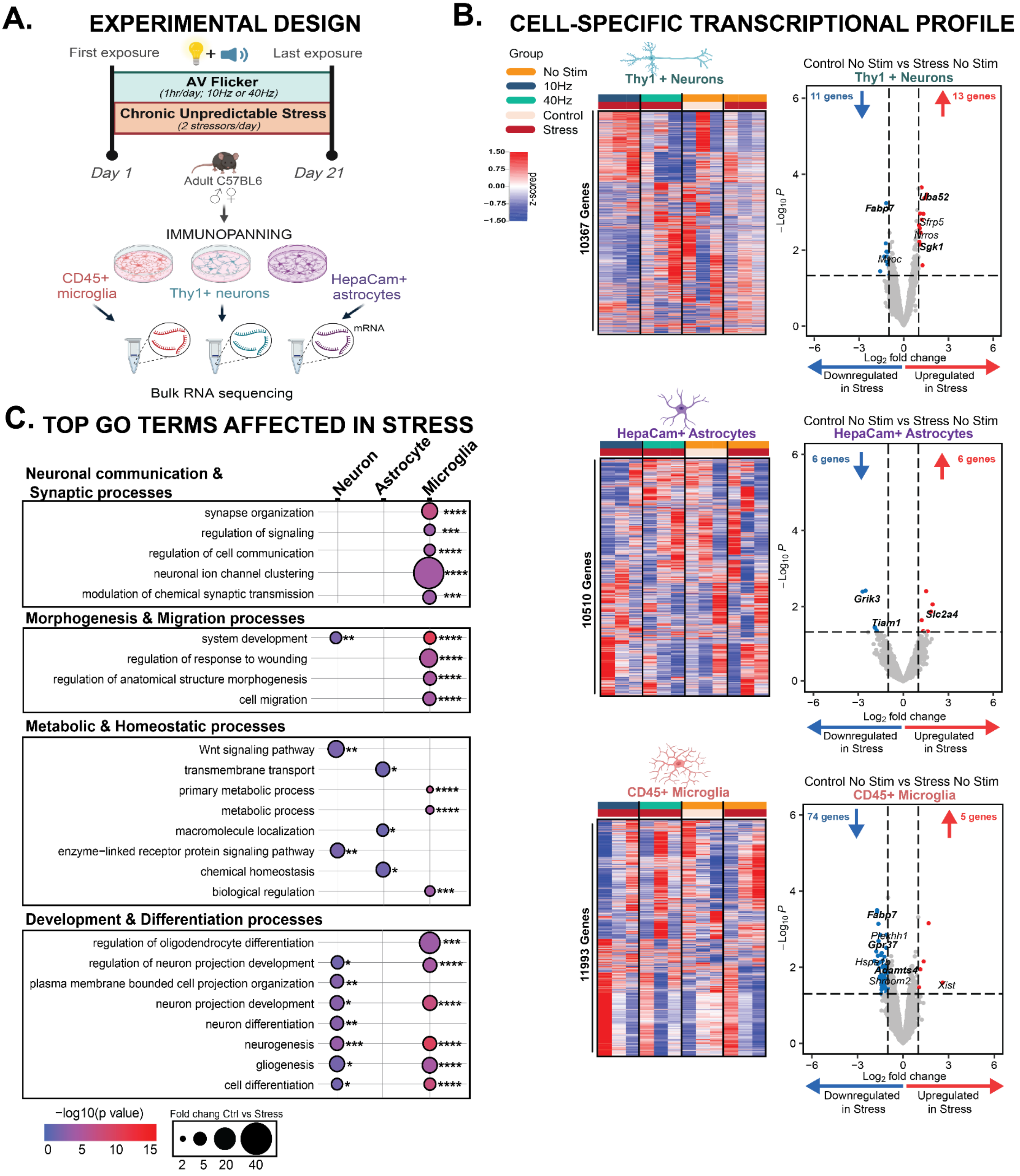


**Supplemental Figure 5: Chronic stress robustly shifts neuronal, astrocyte and microglia transcriptome profile in females. (A)** Experimental paradigm. Thy1+ neurons, HepaCam+ astrocytes and CD45+ microglia were isolated by immunopanning approach within 1 hour of the last stressor exposure on the last paradigm day. **(B)** Heatmaps of gene expression of neurons (top), astrocytes (middle), and microglia (bottom) in control no stim (orange/light pink)

or following stress alone (dark red/orange), stress combined with AV flicker at 10Hz (dark red/blue), or 40Hz (dark red/green). Colors above the heatmap indicate stress (dark red) or no stress (pink) and no stimulation (orange), 10Hz (blue), or 40Hz (green) flicker exposure. In the heatmap, red shows increased expression and blue shows decreased expression compared to no stress no stimulation controls (threshold set. log2fc>1 and $p<0.05$). **(C)** Primary pathways affected by stress alone in neurons, astrocytes, and microglia. n = 3 pooled samples from 6 animals per group. Transcriptional profile at gene expression level unadjusted $p<0.05$, Wald test. GO terms pathway analysis FDR-adjusted *$p < 0.05$, **$p<0.01$, ***$p<0.001$, and ****$p<0.0001$. **A** created in BioRender. Franklin, T. (2025) https://BioRender.com/r17l695

Supplemental Figure 6

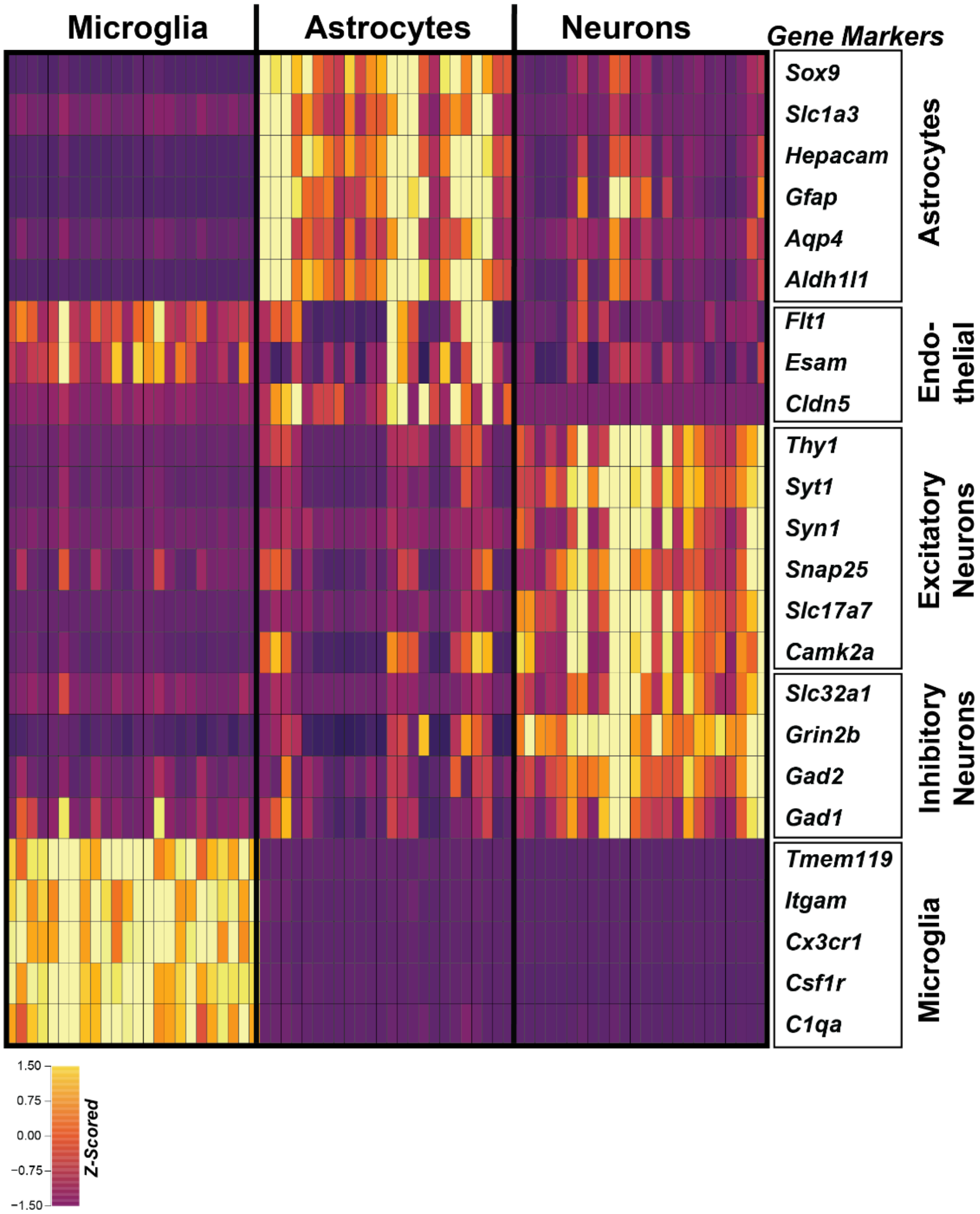


**Supplemental Figure 6: Validation of cell-specific sample enrichment following immunopanning.** Heatmap of cell-type specific gene expression confirming enrichment of neuron-, endothelial-, astrocyte-, and microglia-specific gene sets. Labels at the top indicate samples that are enriched for microglia (left), astrocytes (center), or neurons (right). Yellow indicates high expression, while purple represents low expression. Each row is a cell-type specific gene marker (labeled on the right). Each column is a sample within each cell type. All stress and flicker conditions are included here.

Supplemental Figure 7

#### A. Percentage of Stress Genes Modifiable by AV Flicker

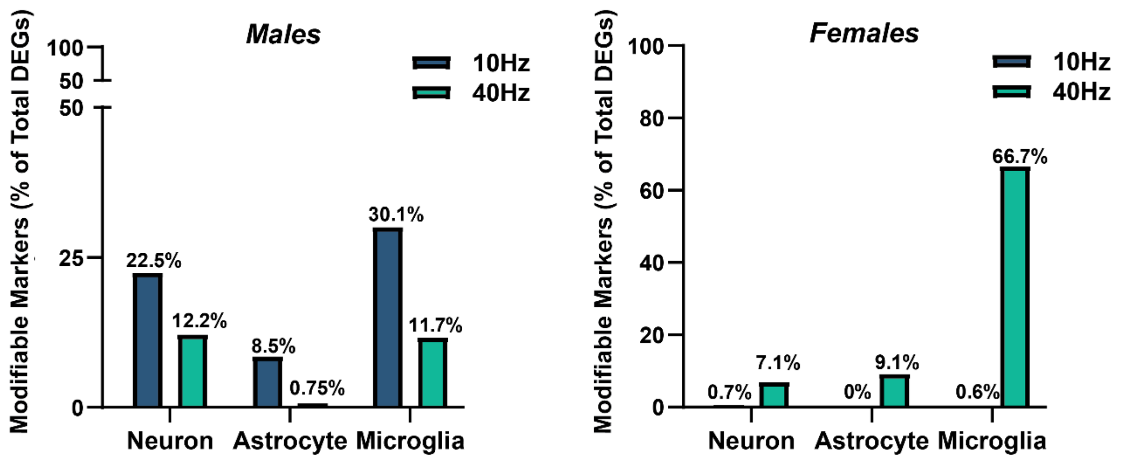


#### B. Shared Modifiable Stress DEGs in AV Flicker Males

| | | |
|---|---|---|
| mt-Nd1 | Cops2 | Scube1 |
| mt-Cytb | Tmem47 | Spock3 |
| Ormdl2 | Tmem126b | Cfap36 |
| Pkia | Mpzl1 | Cdan1 |
| Pcsk7 | Wwp1 | Aspa |
| Cep170 | Ubxn7 | Gm14295 |
| Exoc6 | Fbxo11 | Smim15 |
| Dennd10 | Rbbp9 | Nap1l3 |
| Emc2 | Yju2 | Nae1 |
| Chpt1 | Tial1 | Tspyl5 |
| Ctso | Ralyl | 4930453N24Rik |
| Fgd3 | Lypla1 | 1110059E24Rik |
| Padi2 | | |

#### C. GO Terms from Shared Modifiable Stress DEGs in AV Flicker Males

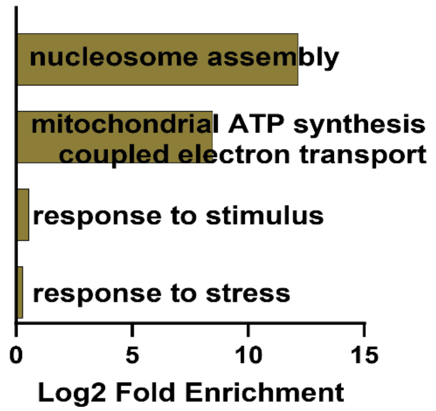


**Supplemental Figure 7: Proportion of AV Flicker-modulated genes that mitigated stress-induced transcriptomic changes across CNS cell types. (A)** Bar graphs showing percent of differentially expressed genes (DEGs) modulated by audiovisual (AV) flicker at 10Hz (grey) or 40Hz (teal) that were classified as "modifiable markers" in stressed male (left) and female (right) mice. Modifiable markers were defined as genes upregulated by stress and downregulated by flicker, or vice versa (see Methods). **(B)** List of shared modifiable DEGs in neurons between 10Hz and 40Hz AV Flicker in stressed male mice. **(C)** Gene ontology (GO) enrichment analysis for overlapping DEGs in neurons from stressed males exposed to 10Hz or 40Hz AV flicker, compared to stressed males with no stimulation. GO terms pathway analysis FDR-adjusted.

Supplemental Figure 8

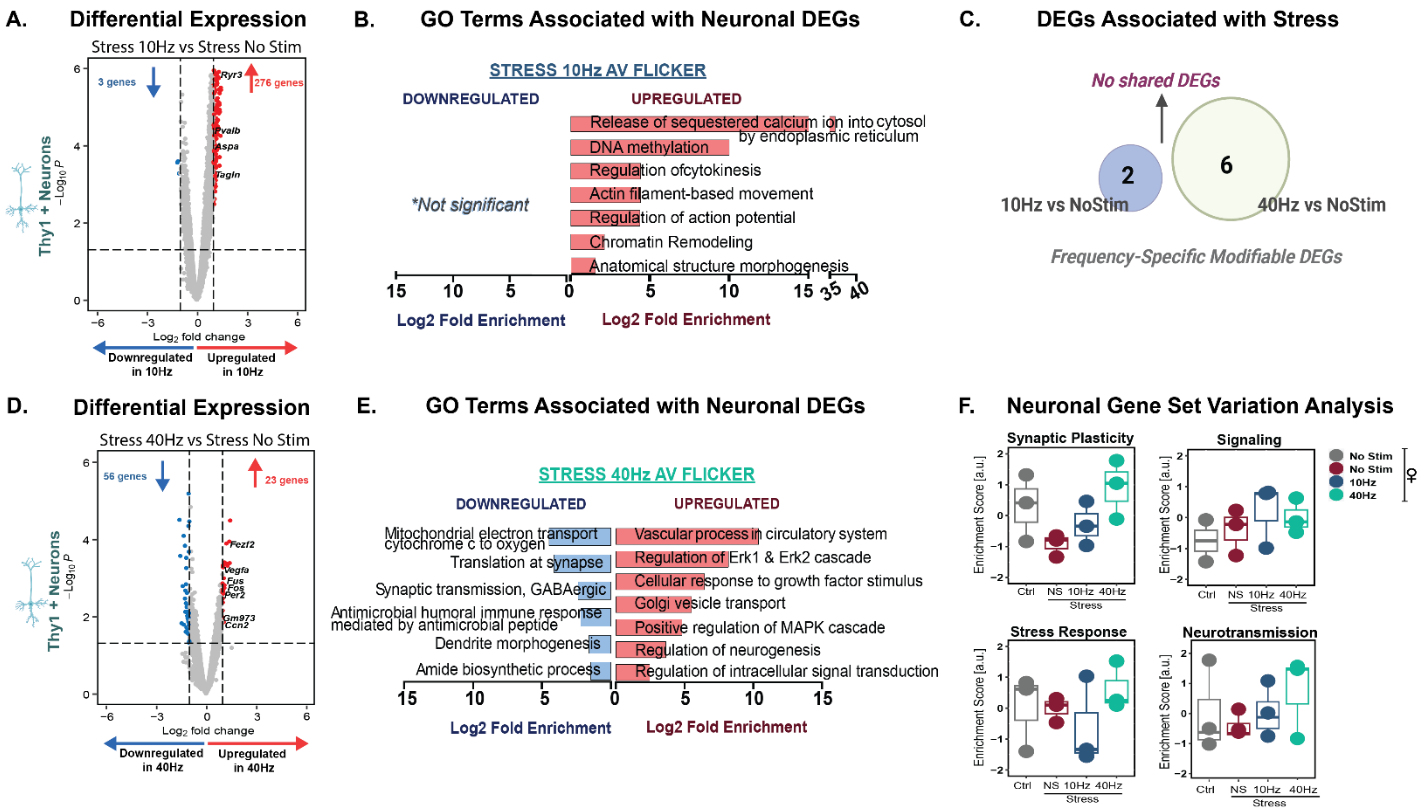


**Supplemental Figure 8: 40Hz AV flicker mitigates stress-induced transcriptional changes in neurons in females. (A)** Volcano plot showing differential gene expression of cortical neurons in stressed females following 10Hz AV flicker compared to stress no stimulation (threshold set. log2fc>1 and p<0.05). **(B)** Gene ontology (GO) enrichment analysis for genes downregulated (blue) and upregulated (red) with 10Hz AV flicker compared to stress no stimulation. **(C)** Venn diagram demonstrating unique (frequency specific) modifiable enriched genes associated with stress following 10Hz (blue) AV flicker compared to 40Hz (green) AV flicker. **(D)** and **(E)** as in A and B for 40Hz AV flicker exposure in stressed females compared to no stimulation stressed females. (**F**) Enrichment scores for specific biological processes (i.e. stress response and synaptic signaling) in stressed female mice different flicker exposure conditions. Box plots show the distribution of scores across samples (median ± interquartile range; n = 3 pooled samples from 6 animals per group). Transcriptional profile at gene expression level without FDR adjustment p<0.05, Wald test. GO terms pathway analysis FDR-adjusted, p < 0.05.

Supplemental Figure 9

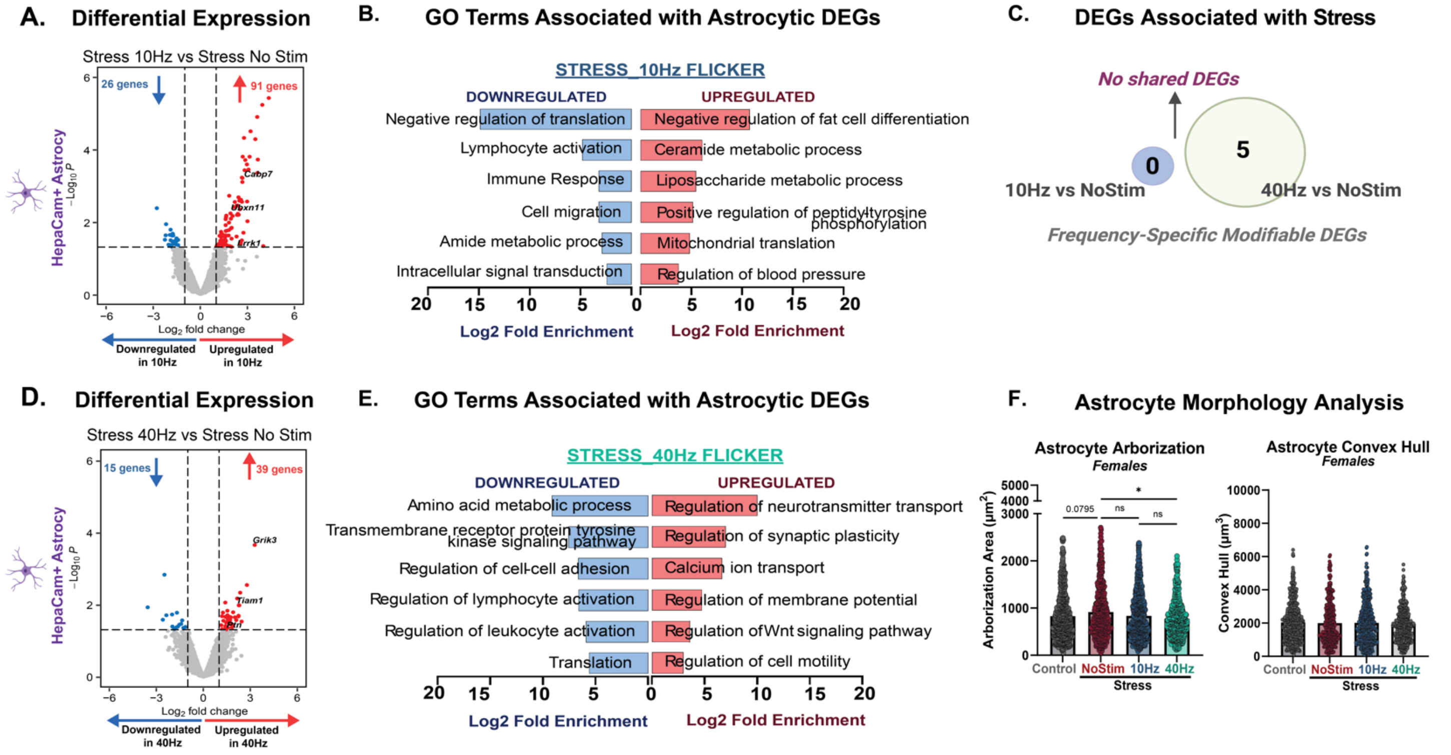


**Supplemental Figure 9: Flicker effects on stress-induced transcriptional changes in frontal cortical astrocytes in females. (A)** Violin plot showing differential gene expression of cortical astrocytes in stressed females following 10Hz AV flicker compared to stress no stimulation (threshold set. log2fc>1 and p<0.05). **(B)** Gene ontology (GO) enrichment analysis for genes downregulated (blue) and upregulated (red) under stress compared to stress no stimulation. **(C)** Venn diagram demonstrating unique modifiable enriched genes associated with stress following (blue) compared to (green). **(D)** and **(E)** as in A and B for 40Hz flicker exposure in stressed females compared to no stimulation stressed females. **(F)** Cortical astrocyte branching (arborization, center), and convex hull volume (right) in female mice after stress and 10Hz AV flicker 10Hz (blue), stress and 40Hz (green), stress alone (red), or no stress no stimulation control (grey) conditions (arborization: H(3) =8.856; p=0.0313; convex hull: H(3) =3.730; p=0.2922). Error bars show mean ± SEM. Each dot is a cell. N = 3 pooled samples from 6 animals per group for transcriptome analysis and n = 431-574 cells from 5 animals per group for morphology analysis. Morphological analyses were conducted at the group level, with data aggregated across cells rather than averaged per subject. Transcriptional profile at gene expression level without FDR adjustment p<0.05, Wald test. GO terms pathway analysis FDR-adjusted, p < 0.05. Kruskal-Wallis test performed for astrocyte morphology analysis followed by Dunn's post hoc test.

Supplemental Figure 10

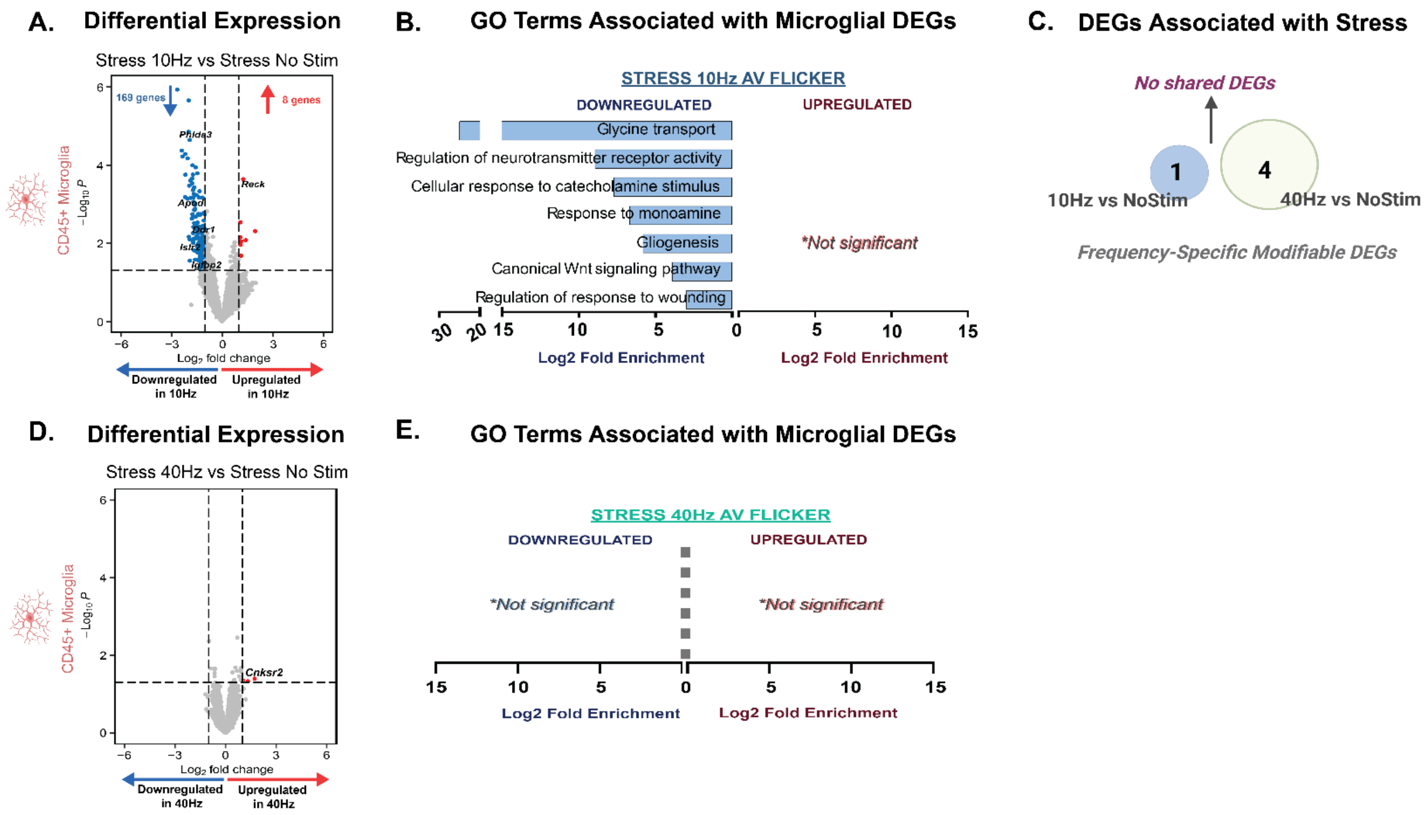


**Supplemental Figure 10: Flicker effects on stress-induced transcriptional changes in frontal cortical microglia in females. (A)** Volcano plot showing differential gene expression in stressed females following 10Hz AV flicker exposure compared to stress no stimulation. **(B)** Gene ontology (GO) enrichment analysis for genes downregulated (blue) and upregulated (red) under after 10Hz AV flicker exposure in stressed females compared to stress no stimulation. **(C)** Venn diagram demonstrating number of unique (frequency specific) modifiable enriched genes associated with stress following 10Hz (blue) AV flicker compared to 40Hz (green) AV flicker. **(D)** and **(E)** as in A and B for 40Hz exposure in stressed females compared to no stimulation stressed males. Note that no GO terms met criteria after 40Hz flicker exposure in microglia of females. N = 3 pooled samples from 6 animals per group for transcriptome analysis. Transcriptional profile at gene expression level without FDR adjustment $p<0.05$, Wald test. GO terms pathway analysis FDR-adjusted $p < 0.05$.

Supplemental Figure 11

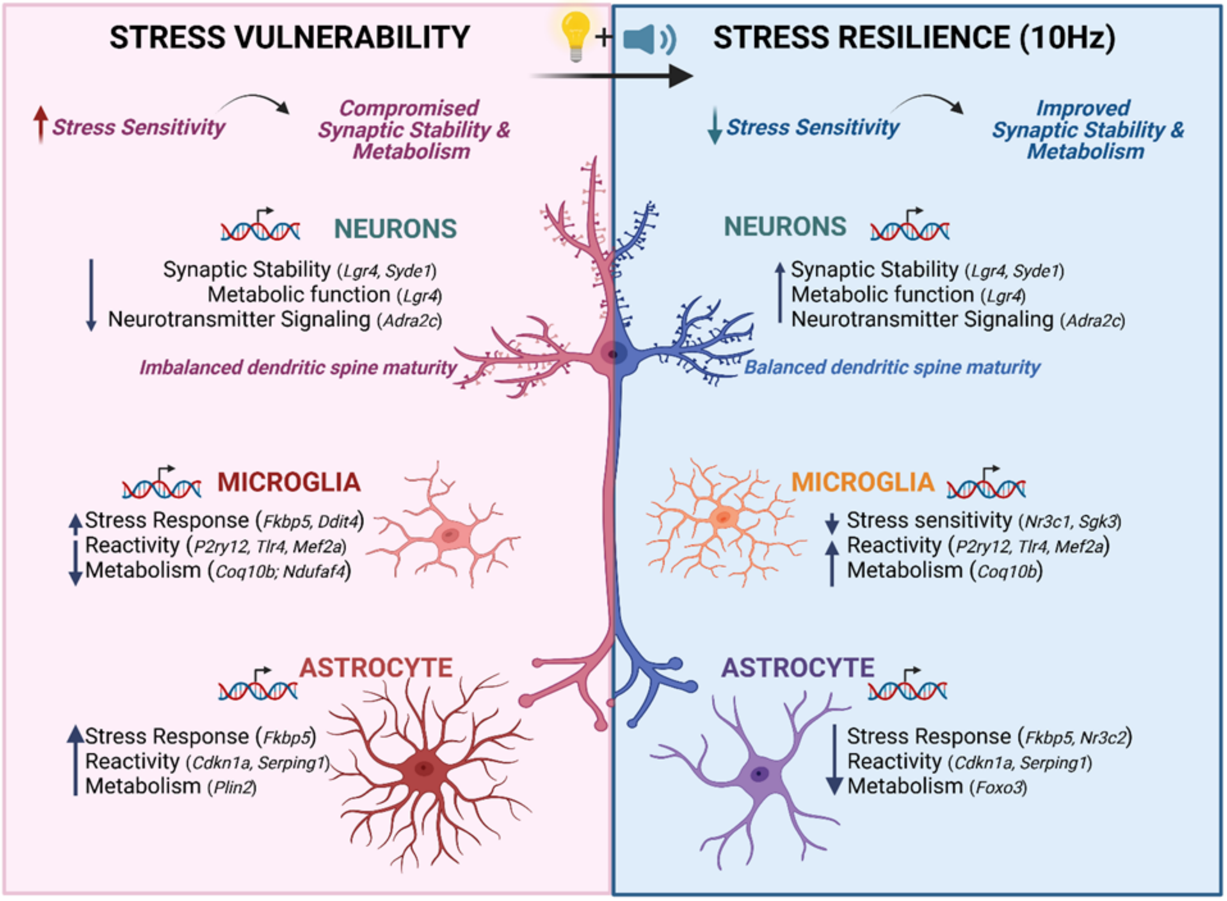


**Supplemental Figure 11: Model of how frequency specific AV flicker mitigates stress pathology through modulation of stress and metabolic signaling.** Frequency-specific AV flicker, particularly at 10Hz in males, mitigates stress-induced pathology by restoring neuronal and glial homeostasis through modulation of synaptic, metabolic, and stress signaling pathways. Chronic stress reduces synaptic stability and compromises metabolic and neurotransmitter signaling, as evidenced by decreased *Lgr4, Syde1,* and *Adra2c* expression among others, which are critical for maintaining synaptic structure and function in cortical neurons. These synaptic alterations, in turn, enhance microglia and astrocyte stress responses and metabolic functions, contributing to neuroinflammation and synaptic dysfunction. 10Hz AV flicker reverses these deficits by enhancing the expression of genes involved in synaptic stability and metabolic function in cortical pyramidal cells, while also modulating the expression of genes involved in the stress responses of microglia and astrocytes. This coordinated response promotes a neuroprotective resilient phenotype. The protective effects of 10Hz AV flicker are further reinforced by robust microglia and astrocyte regulation, accompanied by metabolic regulation across CNS cell types. Stress response adaptations suggest that the resilience in microglia induced by 10Hz AV flicker may promote long-lasting resistance to future stressors, potentially enabling acquired stress resilience over time. Created in BioRender. Franklin, T. (2025) https://BioRender.com/k18t173.

Supplemental Figure 12

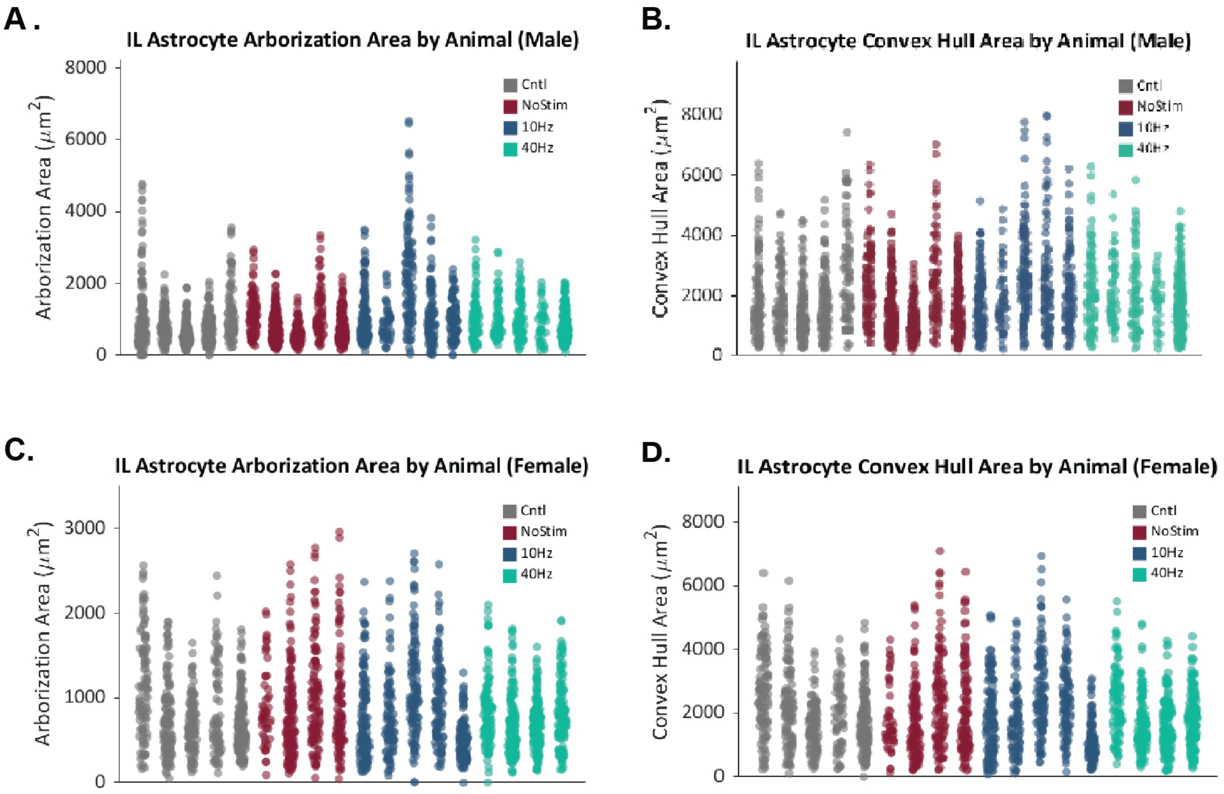


**Supplemental Figure 12: Astrocyte morphology per animal.** (**A**) Individual violin plots per animal (x-axis) showing cortical astrocyte branching (arborization) in male mice that underwent no stress and no stimulation (grey), stress and no stimulation (red), stress and 10Hz flicker (blue), or stress and 40Hz flicker (green) conditions. (**B**) As in A for convex hull volume. (**C**) and (**D**) as in A and B for females.

Supplemental Figure 13

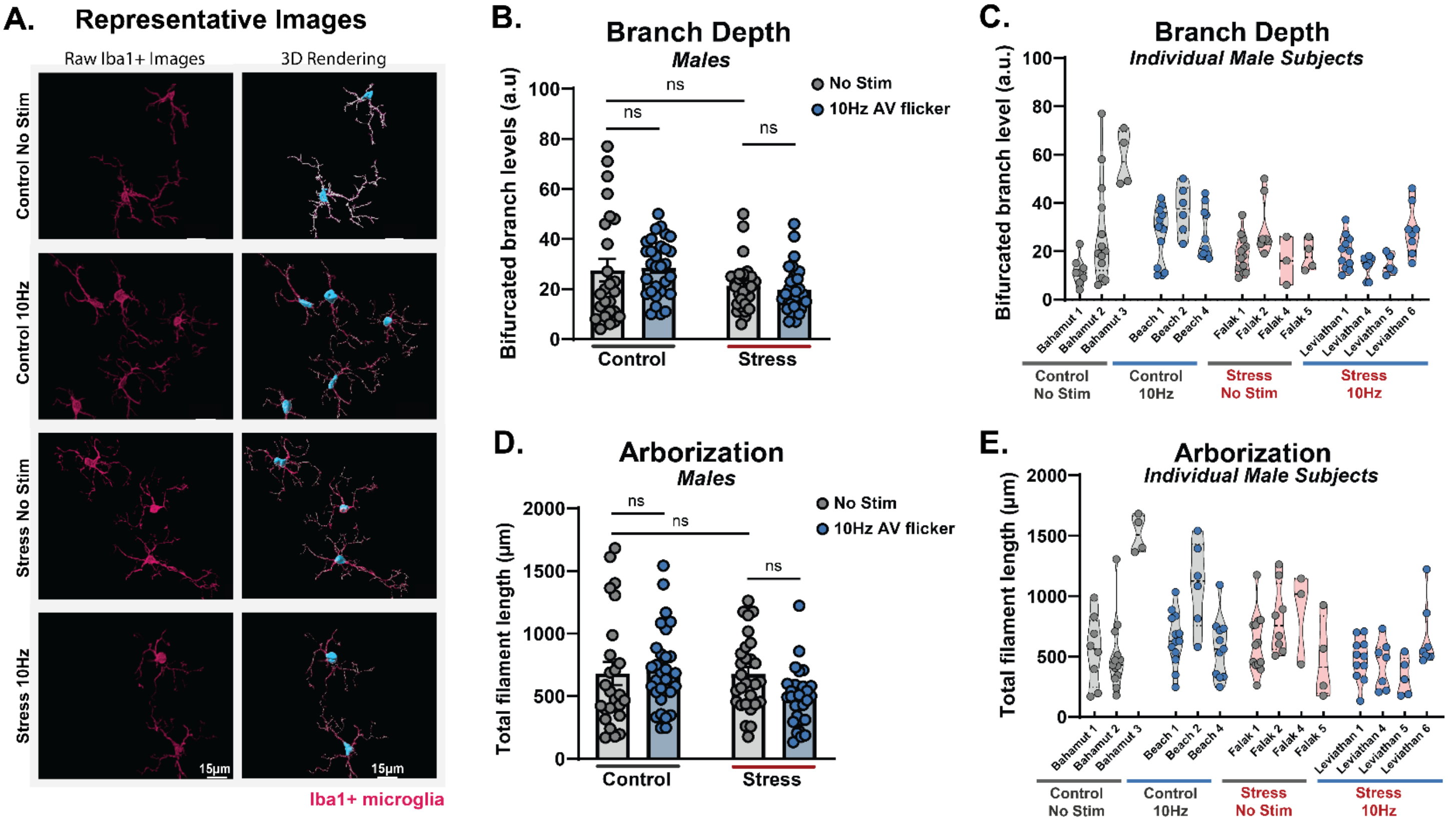


**Supplemental Figure 13: Flicker effects on stress-induced morphological changes in frontal cortical microglia in males. (A)** Example 63x image of Iba1+ microglia (magenta), scale bar 15µm. **(B-C)** Cortical microglia branching depth (bifurcation levels, $F(1, 110) = 8.447$; $p=0.0044$) and **(D-E)** total branching length (arborization, $F(1, 110) = 3.298$; $p=0.0721$) in male mice that underwent no stress and no stimulation (grey bar over grey line), no stress and 10Hz AV flicker (bleu bar over grey line), stress and no flicker (gray bar over red line), or stress and 10Hz flicker (bleu bar over red line) conditions. Error bars show mean ± SEM. Each dot is a cell. N = 25-31 cells from 3-4 animals per group for morphology analysis. Morphological analyses were conducted at the group level, with data aggregated across cells rather than averaged per subject. Two-way ANOVA test performed for microglia morphology analysis followed by Bonferroni's multiple comparisons test.

### Supplementary Tables

**Supplemental Table 1: Neuron DEGs.** Excel table of differentially expressed genes in neurons comparing different stress and flicker conditions. lfcSE: The standard error estimate for the log2 fold change estimate. stat: The value of the test statistic for the gene. pval: p value from Wald test. Padj: Adjusted P-value for multiple testing for the gene.

**Supplemental Table 2: Astrocyte DEGs.** Excel table of differentially expressed genes in astrocytes comparing different stress and flicker conditions. lfcSE: The standard error estimate for the log2 fold change estimate. stat: The value of the test statistic for the gene. pval: p value from Wald test. Padj: Adjusted P-value for multiple testing for the gene.

**Supplemental Table 3: Microglia DEGs.** Excel table of differentially expressed genes in microglia comparing different stress and flicker conditions. lfcSE: The standard error estimate for the log2 fold change estimate. stat: The value of the test statistic for the gene. pval: p value from Wald test. Padj: Adjusted P-value for multiple testing for the gene.

**Supplemental Table 4: GO terms in stress alone in neurons, astrocyte and microglia_unadjusted.** Excel table of gene ontology (GO) biological process in stress alone across neurons, astrocyte and microglia. Count: number of genes associated with specific GO biological process. FE: fold enrichment estimated compared to expected count. pval: p value from Fisher's exact test. Unadjusted (no FDR Correction): unadjusted FDR correction without filtering.

**Supplemental Table 5: Stress associated DEGs modified by 10Hz and 40Hz AV Flicker.** Excel table of differentially expressed genes in stress alone and reversed by flicker conditions across neurons, astrocytes and microglia. lfcSE: The standard error estimate for the log2 fold change estimate. stat: The value of the test statistic for the gene. pval: p value from Wald test. Padj: Adjusted P-value for multiple testing for the gene.

**Supplemental Table 6: Go Terms from shared modifiable stress DEGs in neurons.** Excel table of gene ontology (GO) biological process from differentially expressed genes in stress alone condition and reversed by flicker conditions in neurons. Count: number of genes associated with specific GO biological process. FE: fold enrichment estimated compared to expected count. pval: p value from Fisher's exact test. FDR adjusted: adjusted FDR correction applied.

**Supplemental Table 7: Neuron GO terms_adjusted.** Excel table of gene ontology (GO) biological process from differentially expressed genes in stress alone condition and reversed by flicker conditions in neurons. Count: number of genes associated with specific GO biological process. FE: fold enrichment estimated compared to expected count. pval: p value from Fisher's exact test. FDR adjusted: adjusted FDR correction applied.

**Supplemental Table 8: Astrocyte GO Terms_adjusted.** Excel table of gene ontology (GO) biological process from differentially expressed genes in stress alone condition and reversed by flicker conditions in neurons. Count: number of genes associated with specific GO biological process. FE: fold enrichment estimated compared to expected count. pval: p value from Fisher's exact test. FDR adjusted: adjusted FDR correction applied.

**Supplemental Table 9: Microglia GO Terms_adjusted.** Excel table of gene ontology (GO) biological process from differentially expressed genes in stress alone condition and reversed by flicker conditions in neurons. Count: number of genes associated with specific GO biological process. FE: fold enrichment estimated compared to expected count. pval: p value from Fisher's exact test. FDR adjusted: adjusted FDR correction applied.

**Supplemental Table 10: Key Resources Table.** List of animal strain, materials and software tools used.

**Supplemental Table 11: Statistics.** Statistic reporting for quantified data reported in Figures 1-6 and Supplemental Figures 2-12.
